# Universal structural requirements for maximal robust perfect adaptation in biomolecular networks

**DOI:** 10.1101/2022.02.01.478605

**Authors:** Ankit Gupta, Mustafa Khammash

## Abstract

Consider a biomolecular reaction network that exhibits robust perfect adaptation to disturbances from several parallel sources. The well-known *Internal Model Principle* of control theory suggests that such systems must include a subsystem (called the “internal model”) that is able to recreate the dynamic structure of the disturbances. This requirement poses certain structural constraints on the network which we elaborate in this paper for the scenario where constant-in-time disturbances maximally affect network interactions and there is model uncertainty and possible stochasticity in the dynamics. We prove that these structural constraints are primarily characterized by a simple linear-algebraic stoichiometric condition which remains the same for both deterministic and stochastic descriptions of the dynamics. Our results reveal the essential requirements for maximal robust perfect adaptation in biology, with important implications for both systems and synthetic biology. We exemplify our results through many known examples of robustly adapting networks and we construct new examples of such networks with the aid of our linear-algebraic characterization.

## 1 Introduction

The phenomenon of adaptation is central to biology [1]. Nearly all living systems exhibit some form of adaptation — almost by necessity — as they need to survive and maintain their functions in harsh and unpredictable environments [2]. Also known as homeostasis in biology [3], the property of adaptation refers to the ability of a system to counteract persistent external disturbances and closely maintain some key component (called the “output”) of its internal state. If there is no difference in the steady-state level of this output (called the “set-point”), pre- and post-disturbance, then the system is said to be perfectly adapting. If this stringent form of adaptation holds in spite of perturbations to the parameters or the structure of the network underlying the system, then this property is called robust perfect adaptation (RPA). Biological examples of RPA systems include chemotaxis in prokaryotes [4] and eukaryotes [5], intracellular osmolarity [6], stress response mechanisms [7], transcription factor *σ*-70 activity [8], ammonia uptake [9], copper, zinc, iron and calcium homeostasis [9, 10, 11], glucose uptake [12] and pattern formation mechanisms during development [13].

Evidently the notion of RPA is fairly nuanced and its precise mathematical definition presupposes many specifications, such as how the set-point for the output species is encoded in terms of network parameters, what type of disturbances can affect the network, and how these effects are manifested by the network dynamics. Furthermore this dynamics can have the classical deterministic description or a stochastic description [14] — the latter being more suitable when the network involves biomolecular species with low abundance levels [15]. Existing quantitative studies on RPA systems have mainly focussed on deterministic single-input single-output (SISO) networks where the disturbances can only enter the network dynamics via a pre-specified input node [16, 17, 18, 19]. Even though these SISO networks are quite popular in many engineering domains, the single-input assumption seems unrealistic in biology as disturbances that arise are typically very complex and do not have a single well-defined entry point. In the context of intracellular networks, such disturbances include cross-talk between networks [20], burden effects [21, 22], context dependence [23], unmodelled interactions and a myriad of other extrinsic factors [1]. These disturbances can simultaneously alter several networks parameters that influence the reaction rates, and they can also modify the network structure by adding or removing reaction components. As many of these disturbances cannot be avoided or even mitigated, it makes strong evolutionary sense for intracellular RPA networks to attain a form of *maximal robustness* in which the set-point for the key output variable depends on the least number of network parameters, and it is insensitive to *all* the others. Our aim in this paper is to mathematically characterise and study the structure of biomolecular networks that constitute such maximal RPA (or *maxRPA*) systems.

The scope of our work is very different from existing studies that seek to topologically characterise RPA networks [18, 19]. The primary difference is that we work with a stronger notion of robustness and consequently our characterising conditions for RPA networks are more restrictive than the topological requirements expounded in these studies. In particular our conditions pertain to the stoichiometric structure of a RPA network, and they are both necessary and sufficient for the RPA property to hold and the output set-point to remain invariant to the *largest possible* set of network disturbances. On the other hand, the conditions on the topological structure specified in [18, 19] are only necessary and they only guarantee insensitivity of the output setpoint to the designated input parameter, but this set-point can arbitrarily depend on other network parameters. Moreover, as the topological structure is determined by the signs of the network interactions at the steady-state, it is conceivable that this structure depends non-trivially on the choice of network parameters, further complicating the issue of existence of the RPA property. The other main difference between our work and these existing RPA studies [18, 19] is that in addition to the deterministic setting, we consider the stochastic setting as well. The allows us to delineate the role of dynamical randomness in influencing the RPA requirements.

Motivated by applications in synthetic biology, there have been many exciting developments in recent years in the area of designing biomolecular controller modules that can be interfaced with arbitrary intracellular networks to achieve RPA for some output species of interest. Some of the controller modules that have been proposed and validated, either theoretically and/or experimentally, are the antithetic integral feedback (AIF) controller [8, 24], the autocatalytic controller [25, 26] and the zeroth-order degradation controller [10]. The design paradigm of synthetic biology naturally imposes a clear separation between the controller and the “controlled” networks, which cannot be *a priori* assumed when one studies endogenous RPA networks from the standpoint of systems biology. However a deep result from control theory, called the *internal model principle (IMP)*, stipulates that every RPA network can, at least theoretically, be separated into two components - an *internal model (IM)* that dynamically evaluates the effect of disturbance by measuring the regulated output, and generates counteracting signals, that are then passed to the *rest of the network (RON)* via the actuation mechanism to complete the feedback loop (see Figure 1). Importantly, the IMP states that in order to reject constant-in-time disturbances, the actuation signal computed by IM must measure the time-integral of the *error* signal, which is the difference between the regulated output and its set-point. Hence the IMP should facilitate implementation of the famed *integral feedback* mechanism in control theory that has numerous engineering applications. In the setting of synthetic biology, the controller module (like the AIF, the autocatalytic or the zeroth-order degradation controller) renders the IMP, while the controlled network becomes RON. It is important to point out that IMP does not refer to a single result but rather it provides a template for many possible results; each of which will hold under specific assumptions about the network model and the class of disturbances which are rejected. For example, the IMP has been proved for linear systems [27, 28] and for a certain nonlinear systems where disturbances enter the dynamics linearly [29]. In this paper we provide versions of the IMP in both deterministic and stochastic settings, which are tailored for biomolecular reaction networks exhibiting maxRPA. This allows us to decompose each maxRPA network into an IM and the RON, as in Figure 1, and we explicitly identify an integrator that provides the integral action necessary for robustness.

**Figure 1:**
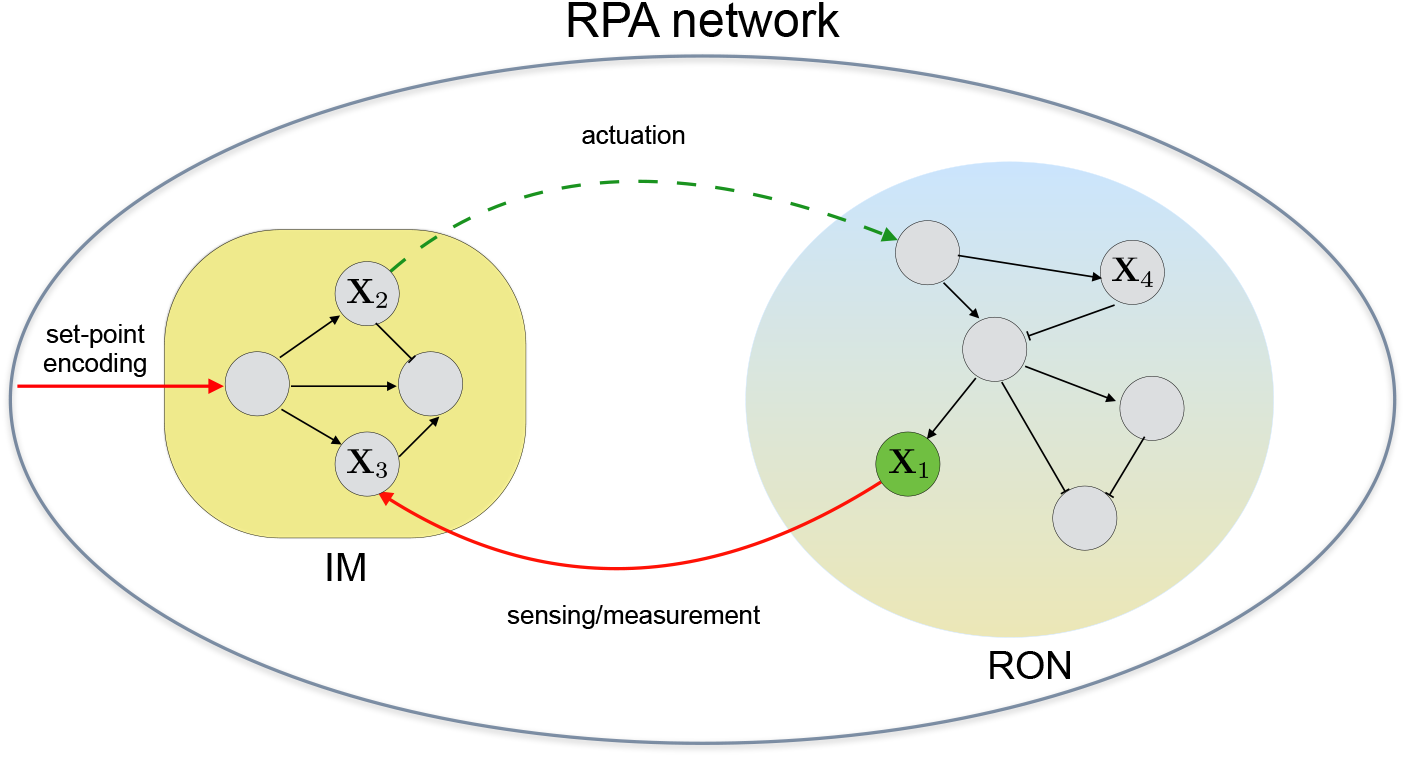
Schematic of the internal model principle (IMP): A RPA network decomposes into an *internal model (IM)* and the *rest of the network (RON)*. Robustness to disturbances is achieved by a feedback mechanism that is realised by the IM. The IM dynamically measures the disturbances via the deviation of the regulated output **X**_1_ from the set-point, and then passes *actuating* signals to the RON to counter the disturbances.

Our main characterisation results posit that maxRPA networks are fully characterised by certain stoichiometric conditions that are simple to state and verify. The principal requirement for maxRPA is a linear-algebraic condition on the network’s stoichiometry, which *remains the same* in both deterministic and stochastic settings (see Sections 3.1 and 3.2). The only difference between the two scenarios is in the requirements for the stoichiometries of the reactions, with the mass-action form [14], whose rate constants are assumed to encode the output set-point. We explicitly identify the set-point encoding function and show that maxRPA characterising conditions are more restrictive in the stochastic case. It follows that stochastic maxRPA networks form a strict subset of deterministic maxRPA networks, which extends a similar result proved in the context of synthetic biology in [24]. Interestingly, our results show that randomness in the dynamics necessitates that the maxRPA network must embed the *antithetic* mechanism, which is characterised by the presence of a sequestration or an annihilation reaction between two IM species (see Section 3.3). We refer to maxRPA networks without such a mechanism as *homothetic* and they can only exist in the deterministic setting. For any maxRPA network we can view its IM as an inbuilt controller that generates robustness for the regulated output variable. The metabolic cost incurred by this controller can be adjudged by the complexity of the IM, in terms of the number of species it contains and the number of reactions among them. It is natural to ask — what are the least complex IMs that are able to confer robustness? We examine this question in both deterministic and stochastic settings and fully characterise the structure of all maxRPA networks with minimal IMs (see Section 4).

In Section 5 we consider known biological examples from the literature and show how they fit our results. These include the bacterial chemotaxis network [4, 30], the network governing proliferative control of cell lineages [31] and the quasi-integral controller based on phosphorylation-dephosphorylation cycles [32]. We also discuss the *sniffer system* [33], which utilises an incoherent feedforward loop (IFFL) to achieve RPA w.r.t. to some of the network parameters, but is not a maxRPA network. Finally in Section 6 we conclude with possible implications of our results.

## 2 Preliminaries

In this section we provide descriptions of reaction networks in both deterministic and stochastic settings. Then we mathematically define our criterion for maxRPA and discuss how this property is connected to the existence of *integrators* [2].

### 2.1 Reaction Networks

Consider a reaction network with *N* biomolecular species **X**_1_,…,**X**_*N*_ that interact among each other through *K* reactions of the form

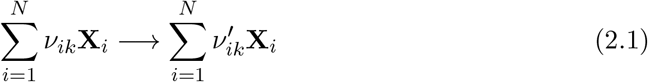

where *ν_ik_* (resp. 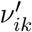) are *stoichiometric* constants in 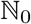 (i.e. the set of nonnegative integers) denoting the number of molecules of species **X**_*i*_ that are consumed (resp. produced) by reaction *k*. Hence the change in the vector of species molecular counts, brought about by a firing of reaction *k*, is given by the *stoichiometric vector* 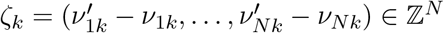 (where 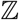 is the set of all integers).

#### Deterministic model

In the classical deterministic model of a reaction network, the state 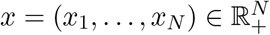 (where 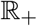 is the set of nonnegative reals) denotes the vector of species concentrations. For each reaction *k*, the state *x* is mapped to the *flux* for reaction *k* according to a *propensity function* λ_*k*_: 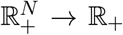. Then the reaction dynamics (*x*(*t*))_*t*≥0_ is described by the following system of ODEs

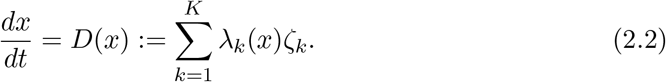

For our results, we shall be assuming that a solution of this system of ODEs exists uniquely for some open set 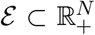 which is time-invariant under the dynamics, and there is a globally attracting fixed point 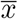, satisfying

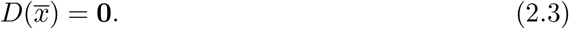

If the propensity function for reaction *k* has *mass-action* form, then it means that λ_*k*_ is given by

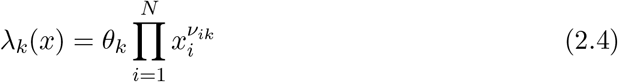

where *θ_k_* is a positive reaction rate constant and *ν_ik_*-s are as in (2.1).

#### Stochastic model

In the standard stochastic model of a reaction network, the state 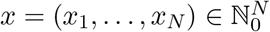 denotes the vector of species copy-numbers or molecular counts. The propensity function λ_*k*_: 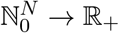 maps the state *x* to the rate of firing λ_*k*_(*x*) of reaction *k*, and when this reaction fires it displaces the state to (*x* + *ζ_k_*) where *ζ_k_* is the stoichiometry vector as before. It is natural to impose the following restriction on each propensity function λ_*k*_

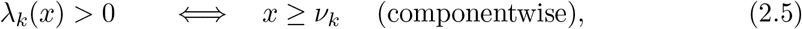

that essentially means that each reaction *k* has a positive rate of firing if and only if the requisite number of reactant molecules (given by *ν_ik_* for species **X**_*i*_) are present. This assumption is biologically reasonable and it ensures that that the non-negative integer orthant 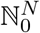 is invariant for the stochastic dynamics. This assumption is trivially satisfied for the propensity function λ_*k*_ if it has the *mass-action* [14] form

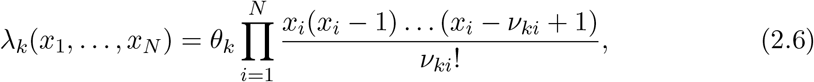

where *θ_k_* is a positive reaction rate constant as before.

In the stochastic model, the reaction dynamics (*X*(*t*))_*t*≥0_ is described by a *continuous-time Markov chain* (CTMC) [14]. Letting 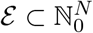 be the set of all accessible states for the CTMC and *p_t_* to be probability distribution of *X*(*t*) i.e.

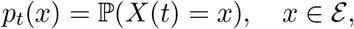

the well-known *Chemical Master Equation* (CME) [14] expresses the time-evolution of *p_t_* as

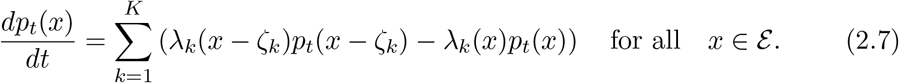

The stability of the CTMC (*X*(*t*))_*t*≥0_ is given by the notion of ergodicity which mandates that regardless of the initial distribution *p*_0_(·), as *t* → ∞, the probability distribution *p*_*t*_(·) converges to a stationary distribution *π* which is a globally attracting fixed point for the CME (2.7). If this convergence is exponentially fast then this property is called exponential ergodicity. In order to verify ergodicity, a two-step approach is needed. First one needs to check if the state-space 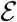, which is typically infinite, is *irreducible*^1^ and then construct a suitable Foster-Lyapunov function^2^ over the state-space 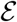 [34]. For stochastic reaction networks, computational frameworks for systematically carrying out both these tasks are provided in [35] and [36] respectively.

### 2.2 The maxRPA property

Suppose that the reaction structure, given by reactions (2.1), is fixed. Henceforth we designate the first species **X**_1_ as the *output* species. The property of RPA pertains to robustly driving the steady-state abundance level of the output species to some set-point *θ*\*, despite *perturbations* or *disturbances* to the network. The steady-state of the network, in both deterministic and stochastic settings, is determined by the propensity functions, which may depend on parameters such as reaction rate constants for mass-action reactions or other parameters, like temperature, which influence the kinetics. It is natural to assume that the set-point is encoded as a function of these parameters, and for definiteness, we suppose that the first *m* reactions in the network have mass-action kinetics and their rate constants *θ*_1_,…, *θ_m_* encode the set-point *θ*\* via some smooth function *ϕ*, i.e.

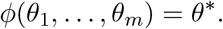

Noting that this set-point will remain invariant to time-rescaling, one can derive a first-order *partial differential equation* (PDE) for *ϕ* which shows that *m* must at least be two (see Section S2 in the Supplement). As we are concerned with maxRPA, we would like our set-point to be determined by the least number of parameters. Hence we set *m* = 2 and in this case our PDE shows that *ϕ* is essentially only a function of the ratio of the parameters *θ*_1_ and *θ*_2_. Denoting this function as *ϕ*_out_, the set-point becomes

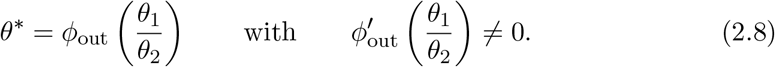

The non-zero derivative condition is added to rule out the trivial case where the setpoint encoding function does not depend on its argument. Our results will show that this set-point encoding function cannot be arbitrary for maxRPA systems and must have a specific form depending on the stoichiometry of the first two reactions.

We shall allow disturbances to enter the system through perturbations of the propensity functions of all the reactions except the first two. These (*K* – 2) reactions need not follow mass-action kinetics and we call the vector of their propensity functions λ = (λ_3_,…, λ_*K*_) as the *uncertain propensity map* which belongs to some pre-defined set Λ_*u*_ of *uncertain* propensity maps. In our setup, constant-in-time disturbances affect the system by perturbing the uncertain propensity map λ to another map in the uncertain set Λ_*u*_.

Given this framework, we now define the maxRPA property in a mathematically precise way. Let *θ* = (*θ*_1_, *θ_2_)* be the vector of rate constants for the first two reactions, which we assume belongs to some feasible open set 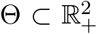, and let *f*: 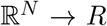 be the projection map, that maps the state vector to the state of the output species

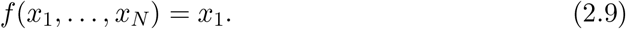

For any fixed (*θ*, λ) ∈ Θ × Λ_*u*_, let (*x*_*θ*,λ_(*t*))_*t*≥0_ be the deterministic dynamics with fixed point 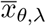. Then we say that the network satisfies the maxRPA property in the *deterministic setting* if and only if

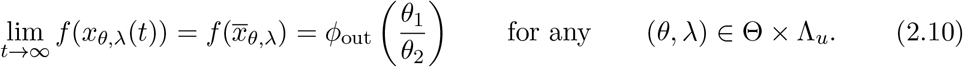

Similarly let (*X*_*θ*,λ_(*t*))_*t*≥0_ be the stochastic dynamics with stationary distribution *π*_*θ*,λ_. Moreover let 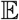 (resp. 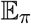) denote the expectation operator under the natural probability distribution (resp. the stationary probability distribution) for CTMC. Then we say that the network satisfies the maxRPA property in the *stochastic setting* if and only if

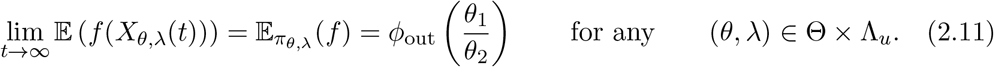

### 2.3 RPA and Integrators

The *internal model principle* (IMP) [27, 29] from control theory implies that for a system to reject constant-in-time disturbances, it must contain a subsystem, called the *internal model* (IM), that is able to generate such disturbances. For this to hold, the subsystem must be an “integrator” as constant disturbances are generated by the ODE

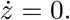

Additionally, IMP stipulates that the internal model IM is only affected by the system output, and it receives no other direct inputs - neither from the rest of the system, nor from the disturbance itself (see Figure 1). Intuitively, IM can be viewed as providing an estimate of the external disturbances, based *only* on the deviation of the output from its set-point.

Consider the deterministic setting and suppose there is a real-valued function *F*(*x*) such that if we let *z*_*θ*,λ_(*t*) = *F*(*x*_*θ*,λ_(*t*)) then we have

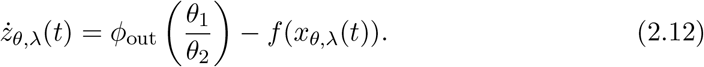

In this case *z*_*θ*,λ_(*t*) is an *integrator* whose time-derivative measures the deviation of the output from the set-point. The function *F*(*x*) is encoded by the IM and it determines the change-of-coordinates that produces the integrator. Observe that for an integrator to achieve its objective of measuring output-deviations, we can allow for more flexibility in the r.h.s. of (2.12). For example, if Ψ(*x*) is any nonlinear function of the state-vector *x* = (*x*_1_,…, *x_N_*) which is monotonic in the first coordinate *x*_1_, then a more general version of the integrator equation (2.12) is given by

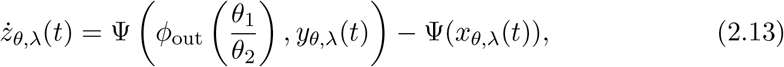

where *y*_*θ*,λ_(*t*) = (*x*_*θ*,λ,2_(*t*),…, *x*_*θ*,λ,*N*_(*t*)) denotes the last (*N* – 1) components of *x*_*θ*,λ_(*t*). This integrator is able to sense the deviation of the output only if the dynamics of *y*_*θ*,λ_(*t*) stays away those values *y* which make the mapping *x*_1_ → Ψ(*x*_1_, *y*) identically zero. Typically Ψ(*x*_1_, *y*) is monomial of variables in *y* and in this case the integrator is often called *constrained* as the dynamics of *y*_*θ*,λ_(*t*) is constrained to stay away from the boundary of 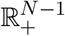 [37].

We now turn to the stochastic setting where the output exhibiting maxRPA is the expectation of the copy-number of species **X**_1_ (see (2.11)). In this case, for a real-valued function *F*(*x*) to specify an integrator we must have that for 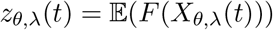

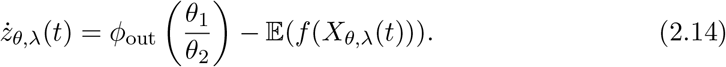

Under the assumption of ergodicity, such a function *F*(*x*) exists uniquely (up to addition by a constant) and it can be found by solving the *Poisson* equation (see Chapter 8 in [38] and [39]). One key difference with the deterministic setting is that we cannot generalise the integrator equation through the introduction of an arbitrary nonlinear function Ψ (see (2.13)), as the expectation operator would not generally commute with the nonlinearity introduced by Ψ. As our results will indicate, this constraint in the form of the integrator equation manifests itself in structural constraints for the maxRPA networks.

We end this section with the simple observation that existence of an integrator is sufficient to guarantee maxRPA in both deterministic and stochastic setting, as the r.h.s. of the integrator equations need to be zero at steady-state.

## 3 Main Results

In this section we present the main results of our paper. These results provide the necessary and sufficient conditions for a network to achieve maxRPA in both deterministic and stochastic settings. The fundamental condition in both cases is a linear-algebraic relation for the stoichiometric structure of the network. Based on this relation we shall classify all maxRPA networks as either *homothetic* or *antithetic* and we shall show that in the stochastic scenario only the latter type of maxRPA networks can arise. We shall also explain how our results provide a novel *internal model principle* (IMP) for maxRPA networks.

Let *S* be the *N* × *K* stoichiometric matrix for the reaction network, defined as

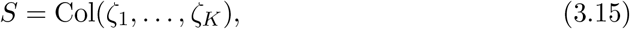

where *ζ_k_* = (*ζ*_1*k*_,…, *ζ_dk_*) is the stoichiometry vector for reaction *k*. This matrix encodes the stoichiometric structure of the reaction network and we shall assume that the set of network reactions is *rich* enough to ensure full row-rank

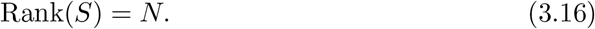

Our principal maxRPA characterising linear-algebraic condition, in both deterministic and stochastic settings, is the existence of a vector *q* = (*q*_1_,…, *q_N_*) and a positive constant *κ* satisfying

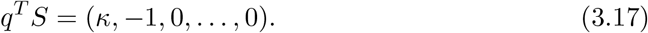

Only the first two components of the r.h.s. vector are nonzero and they are *κ* and −1. For stable networks, if such a pair (*q*, *κ*) exists, then it must be necessarily unique and it can be found by solving a simple linear system of the form *Ax* = *b* (see Lemma S3.1 in the Supplement).

Under certain additional stoichiometric conditions, the structural condition of existence of (*q*, *κ*) satisfying (3.17) characterises all maxRPA systems in both deterministic and stochastic settings. These additional conditions are stricter in the stochastic case, which shows that the class of maxRPA systems in the stochastic scenario is a strict subset of the class of such systems in the deterministic scenario. In the next couple of sections we explicitly state our assumptions and our characterisation results in both deterministic and stochastic settings.

### 3.1 Deterministic maxRPA networks

We now characterise maxRPA networks in the deterministic setting and explicitly identify the form of the set-point encoding function (2.8) as well as an *integrator* that demonstrates the maxRPA property (see Section 2.3). Recall that in our setting, the first two reactions have mass-action propensities with rate constants *θ* = (*θ*_1_, *θ*_2_) ∈ Θ. Hence these propensity functions can be expressed as λ_*k*_(*x*) = *θ_k_m_k_*(*x*) for *k* = 1, 2, where *m_k_* denotes the mass-action monomial

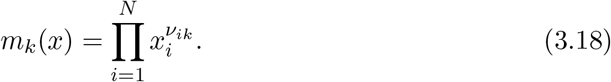

Disturbances enter the reaction system via uncertainties or perturbations in the propensities of the other *K* – 2 reactions. These *K* – 2 propensities constitute the uncertain propensity map λ which belongs to the uncertain set Λ_*u*_. The deterministic reaction dynamics (*x*_*θ*,λ_(*t*))_*t*≥0_ satisfies the following system of ODEs

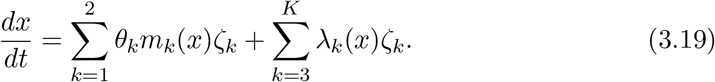

We shall make the following assumptions.

#### Assumption 3.1

*The full row-rank condition* (3.16) *holds and for any fixed* (*θ*, λ) ∈ Θ × Λ_*u*_ *we have the following:*

A. **Stability:** *The ODE system* (3.19) *is well-defined for initial values in some open set* 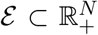 *which is time-invariant under the dynamics and there is a globally attracting fixed point* 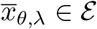.
B. **Local richness of the uncertain set Λ_*u*_:** *There exist R functions ϕ*_1_,…, *ϕ_R_*: 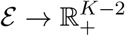 *such that for some ϵ* > 0

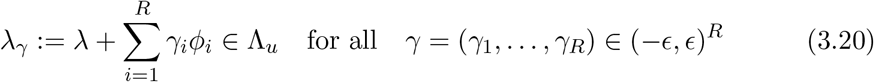

*and if we define a* (*K* – 2) × *R matrix as*

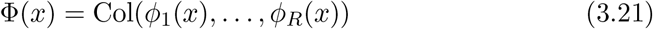

*then this matrix has full row-rank at* 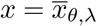.

Part (B) of this assumption implies that around any uncertain propensity map λ, the uncertain set Λ_*u*_ is *locally rich* in the sense that it is possible to solely perturb the propensity for each disturbance inducing reaction *k* ∈ {3,…, *K*} by translating the propensity map λ by a linear combination of functions *ϕ*_1_,…, *ϕ_R_*. Typically we would expect that each propensity gets independently disturbed, i.e. *R* = *K* – 2 and 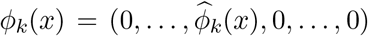. In this case, the rank condition is trivially satisfied and the condition in part (B) reduces to saying that for each *k* = 1,…, *K* – 2, the propensity λ_*k*+2_(*x*) can be perturbed to

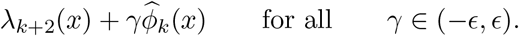

In the special case of mass-action kinetics, i.e.

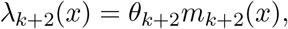

we have

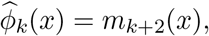

and this condition implies that the disturbance perturbs the rate constant *θ_k_* to a value in (*θ_k_* – *ϵ*, *θ_k_* + *ϵ*). While this covers many scenarios of interest, the general condition, as stated in Assumption 3.1(B), is satisfied by many other forms of disturbances, like those arising from dependent sources like metabolic burden, cross-talk, temperature and other extrinsic factors etc.

Observe that if some component of the uncertain propensity map λ is the zero function, then one cannot perturb it negatively, and hence (3.20) will not hold for all *γ* ∈ (−*ϵ,ϵ*)^*R*^. However our characterisation result will remain valid as its proof will continue to hold upon replacing the usual derivatives w.r.t. components of *γ* with appropriate one-sided derivatives. A zero component in the uncertain propensity map *λ* will naturally arise when one wants to check if adding a disturbance inducing reaction would preserve or violate the maxRPA property.

#### Theorem 3.2

**(Characterisation of deterministic maxRPA networks)** *Suppose that the network reaction dynamics is given by* (3.19) *and Assumption 3.1 holds. Then this network exhibits maxRPA for the output species* **X**_1_ *if and only if the following two stoichiometric conditions are satisfied*:

A. *Reactions* 1 *and* 2 *have as reactants, strictly unequal number of molecules of the output species* **X**_1_ *but equal number of molecules of all other network species, i.e*.

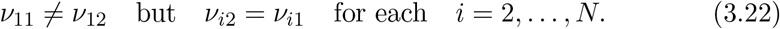
B. *There exists a pair* (*q*, *κ*), *with κ* > 0, *satisfying the linear system* (3.17).

*Moreover, if the network exhibits maxRPA, the set-point encoding function is given by*

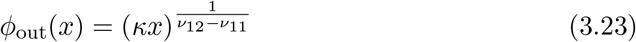

*and an integrator is given by the function*

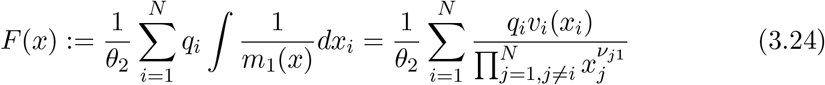

*where*

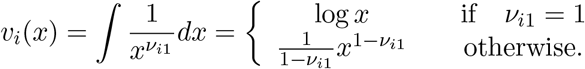

#### Remark 3.3

*Observe that the gradient of integrator* (3.24) *is given by*

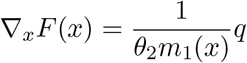

*which shows that if q satisfies the linear system* (3.17) *then for z*_*θ*,λ_(*t*) = *F*(*x*_*θ*,λ_(*t*)) *we have*

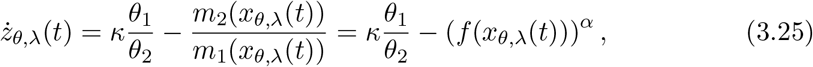

*where α*:= *ν*_12_ – *ν*_11_ *and f*(*x*) = *x*_1_. *The last relation holds because* 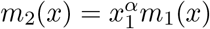 *due to condition* (3.22). *Instead of* (3.24) *we can also define a simpler linear integrator*

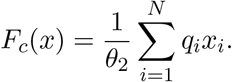

*In this case for z*_*θ*,λ_(*t*) = *F_c_*(*x*_*θ*,λ_(*t*)) *we have*

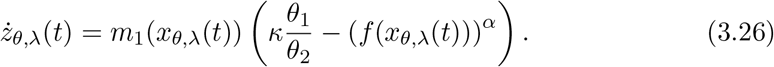

*Both* (3.25) *and* (3.26) *are examples of the general integrator equation* (2.13).

### 3.2 Stochastic maxRPA networks

In this section we provide a result analogous to Theorem 3.2 for maxRPA networks in the stochastic setting. We shall need to make assumptions similar to Assumption 3.1 but these assumptions are more technical in nature in the stochastic setting. We now informally describe these assumptions and their formal description can be found in Section S3.2 of the Supplement.

#### Assumption 3.4

*For any* (*θ*, λ) ∈ Θ × Λ_*u*_ *we have the following:*

A. **Stability:** *The state-space* 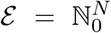 *is irreducible for the CTMC dynamics* (*X*_*θ*,λ_(*t*))_*t*≥0_ *and this CTMC is exponentially ergodic with a unique stationary probability distribution π_θ_*,_λ_.
B. **Richness of the uncertain set** Λ_*u*_: *This criterion is based on the solution of the Poisson equation that defines the integrator* (2.14). *It essentially implies that the set* Λ_*u*_ *of uncertain propensity maps is sufficiently rich to allow perturbations by functions whose linear combination can closely approximate the Poisson equation solution in a certain norm defined using the stationary probability distribution π_θ_*,_λ_.

The full-rank condition (3.16) is necessary for the full nonnegative integer orthant 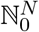 to be an irreducible state-space for the stochastic dynamics (e.g. see [35]). In fact for most biological reaction networks of interest, this condition is also sufficient in ensuring that 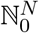 is irreducible. Moreover it has been found that many such networks are indeed exponentially ergodic [36] and this can be shown by constructing a linear Foster-Lyapunov function. The condition on the richness of the uncertain set Λ_*u*_ is more demanding than part (B) of Assumption 3.1, to account for the fact that even at stationarity the dynamics is fluctuating over the whole state-space 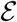.

#### Theorem 3.5

**(Characterisation of stochastic maxRPA networks)** *Suppose that the network reaction dynamics is given by the CTMC* (*X*_*θ*,λ_(*t*))_*t*≥0_ *and also suppose that Assumption 3.4 holds. Then this network exhibits maxRPA for the output species* **X**_1_ *if and only if the following two stoichiometric conditions are satisfied:*

A. *Reactions* 1 *and* 2 **do not have** *molecules of species* **X**_2_,…, **X**_*N*_ *as reactants, and* **exactly one** *of them involves a* **single** *molecule of the output species* **X**_1_ *as reactant while the other involves* **none** *i.e*.

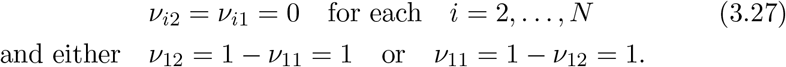
B. *There exists a pair* (*q*, *κ*), *with κ* > 0, *satisfying the linear system* (3.17).

*Moreover, if the network exhibits maxRPA, the set-point encoding function is given by*

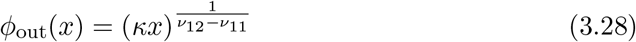

*and an integrator is given by the linear function*

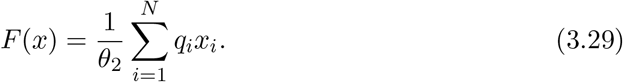

#### Remark 3.6

*Observe that the stoichiometric conditions imposed by this theorem are stricter that those imposed by Theorem 3.2. This shows that if the underlying assumptions hold, then the set of networks exhibiting maxRPA in the stochastic setting is a strict subset of the set of networks exhibiting maxRPA in the deterministic setting. Also note that in this stochastic case the difference in the number of reactant molecules for the output species* **X**_1_, *between reactions 1 and 2 is either* 1 *or* −1. *Hence ν*_12_ – *ν*_11_ ∈ {−1,1} *and so the set-point encoding function can also be expressed as*

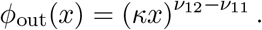

### 3.3 Homothetic and Antithetic maxRPA networks

In the previous sections, we characterised networks achieving maxRPA in both deterministic and stochastic settings. These networks comprise *N* species **X**_1_,…,**X**_*N*_ and from now on we add the null species 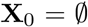 in our species-set and we adopt the convention that

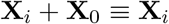

for any species **X**_*i*_. Our key stoichiometric criterion for RPA characterisation, which is common to both deterministic and stochastic settings, is the existence of a unique solution (*q*, *κ*) to the linear-algebraic system (3.17). Letting *q* = (*q*_1_,…, *q_N_*), we can think of *q_i_* as the *charge* of species **X**_*i*_ and to the null species **X**_0_ we assign the charge of *q*_0_ = 0. Based on these charge values, we can partition the species set 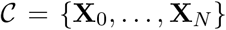 of a maxRPA network into the set of positively charges species 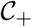, negatively charged species 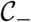 and neutral species 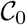

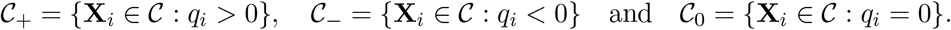

Based on this partition we classify all maxRPA networks as follows:

#### Definition 3.7

**(Classification of maxRPA networks)** *A maxRPA network is called* **homothetic** *if either* 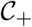 *or* 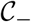 *is empty. On the other hand, a maxRPA network is called* **antithetic** *if both* 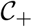 *and* 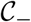 *are nonempty*.

In other words, for a maxRPA network to be homothetic, all the nonzero entries in vector *q* must have the same sign, while if *q* has least two nonzero entries with opposite signs, then this network is antithetic.

The linear-algebraic condition (3.17) implies that for each disturbance inducing reaction *k* ∈ {3,…, *K*}, its stoichiometric vector *ζ_k_* is orthogonal to q, i.e. *q^T^ζ_k_* = 0. Hence each of these reactions are *charge-preserving* in the sense that the cumulative charge of the reactant complex is the same as that of the product complex

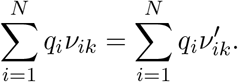

Consider the common scenario where all reactions are *bimolecular*, i.e. all reactions can have at most two reactants i.e. 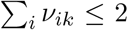. In this scenario antithetic RPA networks are marked by the presence of a generalised *sequestration* or *annihilation* reaction of the form

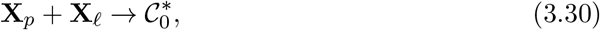

where 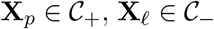 and 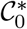 denotes any combination of species that belong to the set 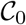 (see Remark S3.5 in the Supplement). Note that such a reaction cannot arise in a homothetic network because either 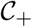 or 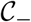 is empty.

### 3.4 A novel *Internal model principle* (IMP)

We now discuss how our characterisation results provide a novel IMP for maxRPA networks, as illustrated in Figure 1. Let 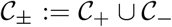 denote the set of species that form the support for vector *q* i.e. *q_i_* ≠ 0 if and only if 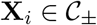. We can view the species in 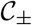 as forming the *internal model* (IM) that generates external disturbances via the integrator (see Section 2.3) defined by function *F*(*x*) identified in Theorems 3.2 and 3.5.

Typically, one is interested in situations where the disturbance inducing reactions are able to independently perturb the output species **X**_1_

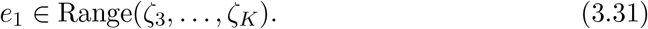

In particular, if there exists a reaction that creates or degrades a molecule of **X**_1_, then this condition is trivially satisfied. An example where this condition does not hold is the simple birth-death module

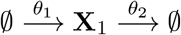

which may include several disturbance inducing reactions that do not alter the copynumbers of the output species **X**_1_. In this example the IM includes the output species **X**_1_ and the property of RPA is trivial because the output is unaffected by the disturbances. On the other hand, if condition (3.31) holds then the IM cannot contain the output species **X**_1_ (see Lemma S3.2 in the Supplement) and for it to perform integral action and generate robustness, there must exist one or more actuation reactions from the IM to the rest of the network (RON). These actuation reactions complete the feedback loop and their presence is necessary for the stability of the full RPA network (see Figure 1). Generally actuation reactions are a subset of disturbance inducing reactions, but it is possible for reaction 1 and/or 2 to serve as actuation reactions (see Section 5.1).

### 3.5 All stochastic maxRPA networks are Antithetic

Our next result shows that in the stochastic setting any maxRPA network must embed the so-called *antithetic* motif, which was first discovered in [8], thereby showing that the antithetic topology is “universal” in the stochastic setting. This result extends a similar universality result proved in [24], for the situation inspired by synthetic biology applications, where the controller network is separate from the network being controlled, and the latter is arbitrary. We shall assume that the network is bimolecular, and as mentioned in Section 3.3 a key property of antithetic networks is the existence of a generalised annihilation or sequestration reaction of the form (3.30) between the IM (comprising species in 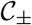) and the RON (comprising species in 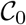).

#### Theorem 3.8

**(Generic decomposition of stochastic maxRPA networks)** *Suppose that Assumption 3.4 and condition* (3.31) *holds and all reactions are bimolecular. Then for a maxRPA network in the stochastic setting we have the following up to renaming of species* **X**_2_,…, **X**_*N*_ *and interchanging of reactions 1 and 2:*

A. *The maxRPA network must be antithetic*.
B. *There is a species* 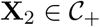 *such that the first reaction is of the form*

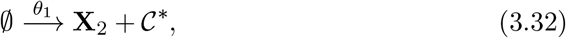

*where* 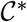 *denotes any combination of network species*.
C. *There is a species* 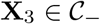, *such that the second reaction is of the form*

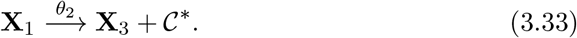
D. *There exists at least one generalised sequestration reaction of the form* (3.30). *Moreover reactions of this form are the only possible reactions where the products are species in* 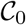 *and one of the reactant species is in* 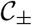.

#### Remark 3.9

*We refer to* (3.32) *as the set-point encoding reaction (as its reaction rate θ*_1_ *sets the numerator of the set-point) and to* (3.33) *as the output sensing reaction (as its reaction rate is θ*_2_*x*_1_ *and x*_1_ *is the output species copy-number). The encoding and sensing is performed by the internal model (IM) (comprising species in* 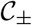) *via production of distinct species* **X**_2_ *and* **X**_3_ *(the species are renamed if necessary) in the IM. The generic decomposition of stochastic maxRPA networks is depicted in Figure 2*.

**Figure 2:**
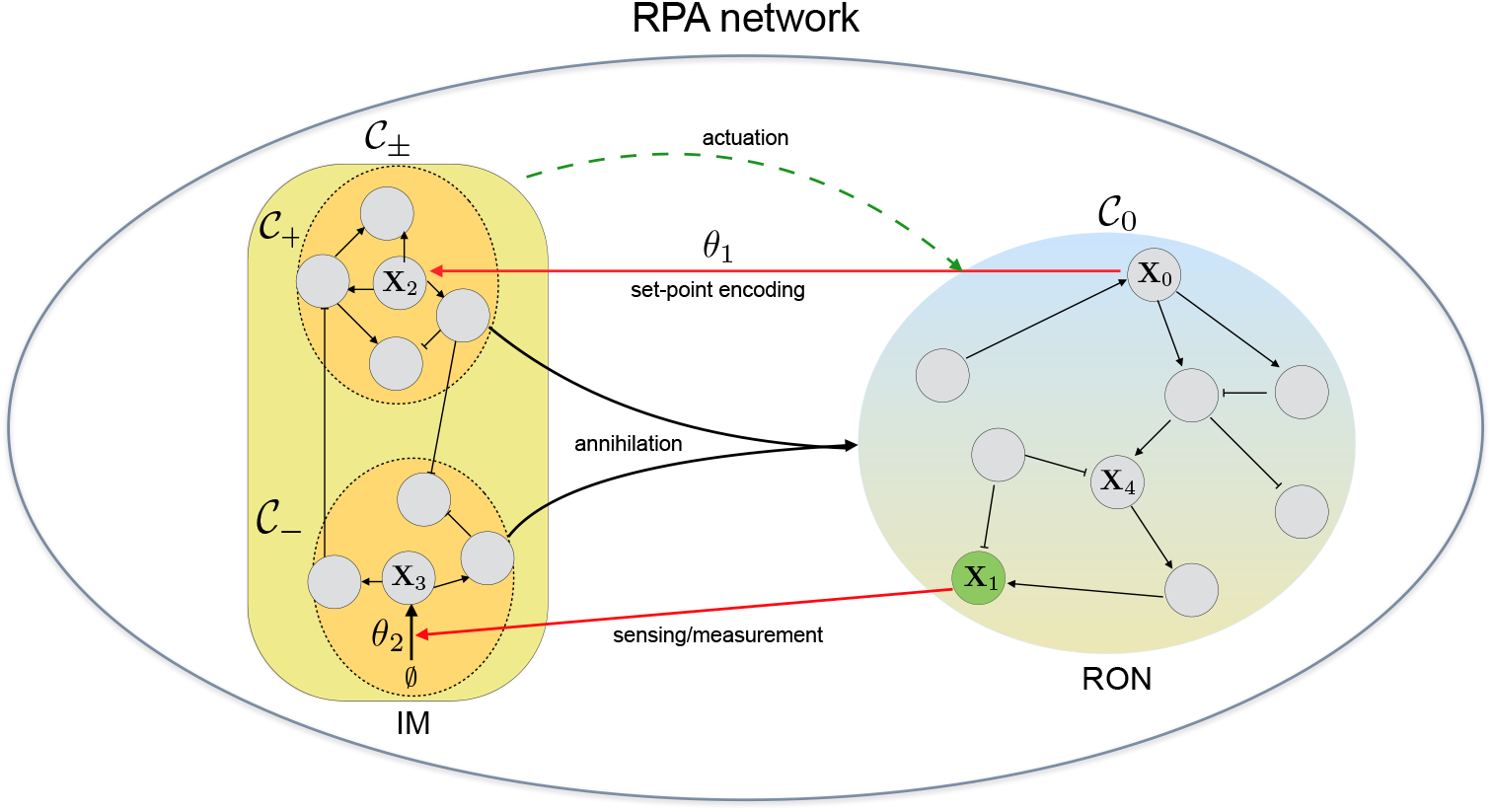
Generic decomposition of stochastic maxRPA networks. Depicts the generic structure of a maxRPA network in the stochastic setting, as proved by Theorem 3.8. Such a network decomposes into an *internal model* (IM), comprising species in 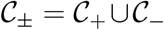, and the *rest of the network* (RON) with species in 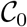. There is a set-point encoding reaction (3.32) producing species 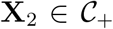 and a sensing reaction (3.33) that measures the output species **X**_1_ via production of species 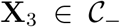. All reactions with reactants in IM and products in RON must have the generalised annihilation form (3.30) and there must exist at least one such reaction, as indicated.

## 4 maxRPA networks with minimal IMs

The internal model (IM) of a maxRPA network characterises the embedded control mechanism that generates the RPA property. In this section we identify the minimal IM necessary for maxRPA where by minimality we mean the IM with the least number of species and the least number of reactions, in that order. This minimality question has important consequences from both systems and synthetic biology perspectives. For example, one might surmise that evolution will seek out the least complex designs for the IM module as they would bestow maxRPA with the smallest amount of extra metabolic burden [21, 22]. This also makes minimal complexity an especially useful design criterion in synthetic biology where building complex circuits is fraught with many difficulties like loading effects [40], cross-talk [20] etc.

Of course the least complex IM would be the one which has only one species. We call this a *singleton* IM and in Section 4.1 we explore the structure of maxRPA networks with such IMs. Observe that these networks must be *homothetic* and the dynamics must be deterministic as in the stochastic setting any maxRPA network must necessarily be antithetic and its IM should comprise least two species (see Theorem 3.8). Hence networks with singleton IM only exhibit maxRPA in the deterministic setting and not the stochastic setting, illustrating that the containment of the set of stochastic maxRPA networks within the set of deterministic maxRPA networks is strict (see Remark 3.6). The least complex IM for an antithetic maxRPA network would consist of exactly two species and we refer to such an IM as *dyadic*. In Section 4.2 we explore the structure of maxRPA networks with dyadic IMs.

### 4.1 Homothetic networks with *singleton* IMs

Consider a deterministic maxRPA network whose IM consists of only the species **X**_2_, i.e. 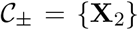. By definition this network is homothetic and we shall assume that condition (3.31) holds. Let (*q*, *κ*) be the maxRPA characterising pair that satisfies (3.17). Then only the second component of *q* (i.e. *q*_2_) is non-zero while the rest are all zeros. From (3.17) we can conclude that the copy-numbers of **X**_2_ are not altered by the disturbance inducing reactions (i.e. *ζ*_2*k*_ = 0 for *k* = 3,…, *K*) and for the first two reactions we have

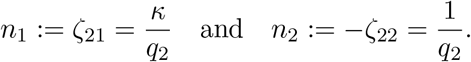

Recall that the nonnegative integer *ν_ik_* denotes the number of molecules of species **X**_*i*_ that are consumed as reactants by the *k*-th reaction. Due to Theorem 3.2 the first two reactions, with mass-action kinetics, must have the form

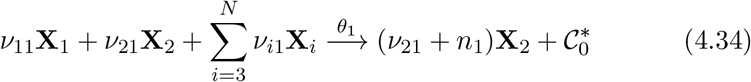

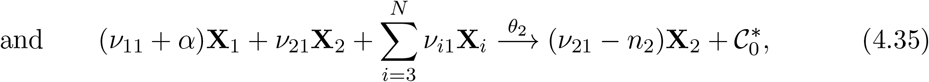

where *α* is some non-zero integer satisfying *α* ≥ – *ν*_11_ and 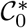 denotes any combination of species in 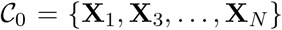. Recall from Remark 3.9, that reaction 1 is the set-point encoding reaction, while reaction 2 is the output sensing reaction. Note that since *κ* > 0, both *n*_1_ and *n*_2_ must have the same sign as *q*_2_ and the output set-point encoding is given by

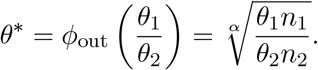

#### 4.1.1 Zeroth-order degradation networks

Consider the case *ν*_21_ = 0 in which the lone IM species **X**_2_ is not a reactant in either the first reaction (4.34) or the second reaction (4.35). This implies that exactly one of these two reactions causes *negative production* or loss of **X**_2_ molecules, violating the reaction form (2.1). The rate of this reaction is not proportional to the abundance of **X**2 molecules which can lead to negative abundance levels for this species. Such a reaction is called *zeroth-order degradation* and even though it is non-physical it can be approximately realised when the degradation is assisted by an enzyme which operates at saturation [10].

Depending on whether *q*_2_ is positive (i.e. 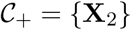) or negative (i.e. 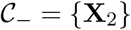), two types of maxRPA networks based on zeroth-order degradation can be constructed. An example of both these network types is shown in Figure 3. In both the examples, we set *ν*_*i*1_ = 0 for all *i*. The key difference between them is that *q*_2_ is positive in one case (panel A) while it is negative in the other (panel B). In the former, the zeroth-order degradation reaction acts like a sensing reaction while in the latter, it acts like a set-point encoding reaction.

**Figure 3:**
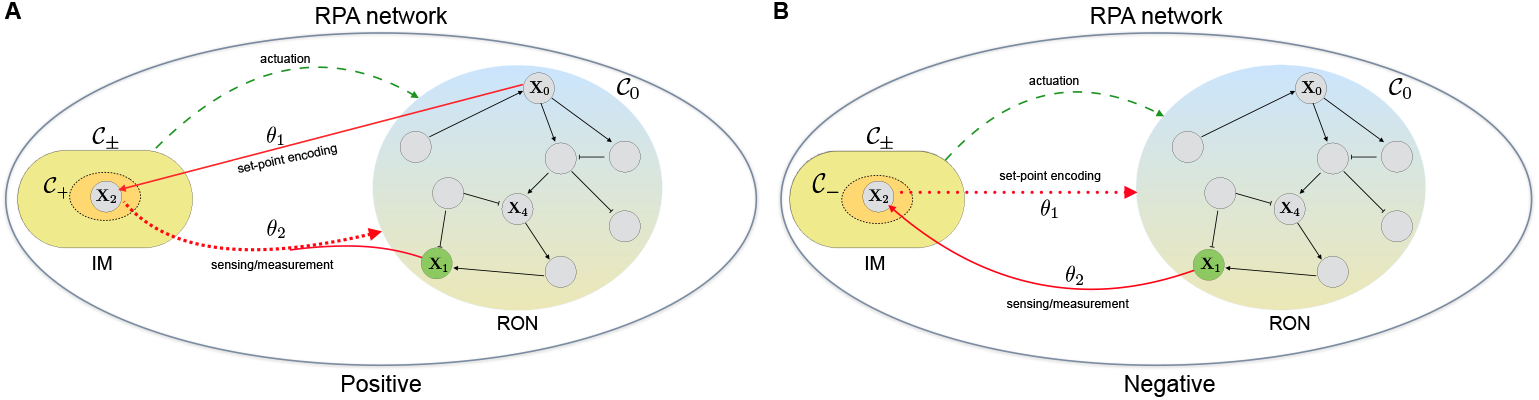
Deterministic maxRPA networks with singleton internal models (homothetic) and zeroth-order degradation. The internal model (IM) consists of only one species 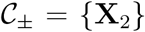 whose charge *q*_2_ is either positive (panel A) or negative (panel B). All the other species belong to the rest of the network (RON). The zeroth-order degradation reaction is shown with a dotted arrow.

#### 4.1.2 Autocatalytic networks

Now consider the case *ν*_21_ > 0 in which the lone IM species **X**_2_ is a reactant in both the first reaction (4.34) and the second reaction (4.35). Hence, irrespective of the sign of *q*_2_, **X**_2_ stimulates its own production and therefore we call such networks *autocatalytic*.

In Figure 4 we present two autocatalytic maxRPA networks that resemble the networks in Figure 3, except that non-physical zeroth-order degradation reactions are replaced by autocatalytic reactions. In both the examples we set *ν*_*i*1_ = 0 for all i except *i* = 2. As before, the difference between them is that *q*_2_ is positive in one case (panel A) while it is negative in the other (panel B). In the former, we choose *ν*_21_ = *n*_2_ and so reaction 1 (set-point encoding) is autocatalytic, while in the latter we choose *ν*_21_ = −*n*_1_ and so reaction 2 (output sensing) is autocatalytic.

**Figure 4:**
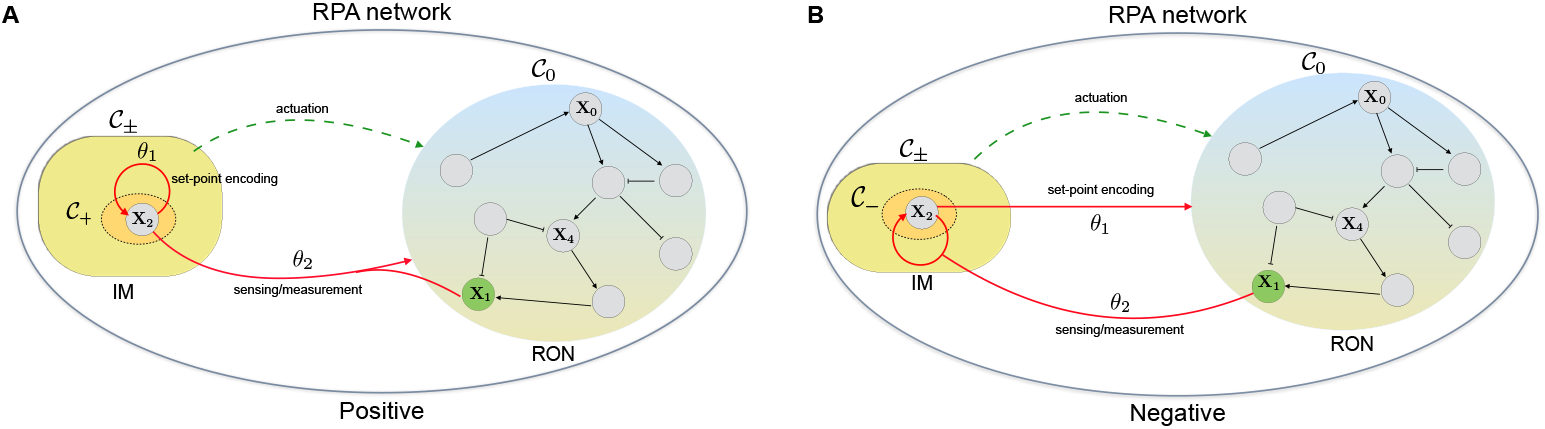
Deterministic maxRPA networks with singleton internal models (homothetic) and autocatalytic reactions. The internal model (IM) consists of only one species 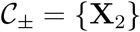 whose charge *q*_2_ is either positive (panel A) or negative (panel B). All the other species belong to the rest of the network (RON). The autocatalytic reaction is shown by a loop.

In the special case of *ν*_21_ = −*n*_1_ = *n*_2_ = 1, 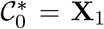 in (4.34) and 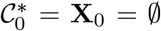 in (4.35), these two reactions encode the simplest network exhibiting the property of *Absolute Concentration Robustness* (ACR), whereby one of the species (**X**_1_ in our case) has the same value at any equilibrium value. For more details we refer the readers to the celebrated paper of Shinar and Feinberg [41] which present sufficient structural conditions for networks to exhibit ACR. In a recent paper the existence of a linear *constrained* integrator for a wide class of ACR networks is shown [42].

### 4.2 Antithetic maxRPA networks with *dyadic* IMs

Consider an antithetic maxRPA network in the deterministic setting whose IM consists of two species **X**_2_ and **X**_3_, i.e. 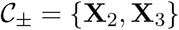. Let (*q*, *κ*) be the maxRPA characterising pair that satisfies (3.17). Without loss of generality we can assume that *q*_2_ > 0 (i.e. 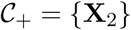) and *q*_3_ < 0 (i.e. 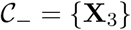). All the other components of the *q* vector are zero. In this section we shall assume that all reactions are *bimolecular* i.e. 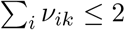 for each reaction *k* of the form (2.1).

From (3.17) we can conclude that for each disturbance inducing reaction *k* = 3,…, *K* we must have

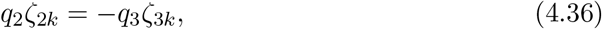

and so *ζ*_2*k*_ and *ζ*_3*k*_ have the same sign. Hence each disturbance inducing reaction either simultaneously *degrades* or *produces* both species **X**_2_ and **X**_3_. Suppose that one such reaction *k* causes degradation of both the species, then as this reaction is bimolecular it must be of the generalised sequestration (3.30) form

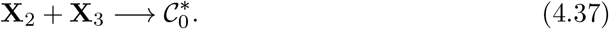

Since *ζ*_2*k*_ = *ζ*_3*k*_ = −1, due to (4.36) we have *q*_3_ = −*q*_2_. To preserve dynamical stability, such a degradation causing reaction must indeed exist provided one of species **X**_2_ or **X**_3_ is only produced by the first two reactions, and not degraded.

Let us examine the form of the first two reactions more closely under the restrictions imposed by Theorem 3.2. Without loss of generality we can assume that *α* > 0, as the other case is symmetric. As all reactions are bimolecular, there must exist a species **X**_*m*_ such that the first two reactions, with mass-action kinetics, have the form

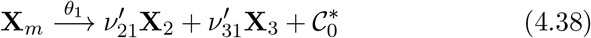

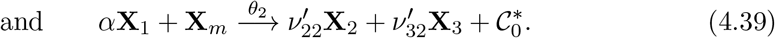

Here **X**_*m*_ can be any species, including the null species 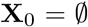 for *m* = 0. Note that for the second reaction to be bimolecular, *α* can be 2 only when *m* = 0, and for other values of *m*, *α* should be 1. We now separate our analysis into two cases, depending on the value of *m*.

**Case 1:** *m* ∉ {2, 3}: In this case *q_m_* = 0 and (3.17) implies that

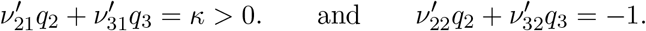

Since *q*_2_ > 0 and *q*_3_ < 0 we must have that 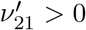 and 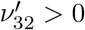 which says that the first reaction must produce species **X**_2_ while the second reaction must produce species **X**_3_. As these two reactions are not degrading any **X**_2_ or **X**_3_, there must exist a disturbance inducing reaction of the form (4.37), as we explained earlier, which in turn implies that *q*_3_ = −*q*_2_. We illustrate this case with an example shown in Figure 5 where we set 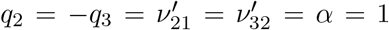 and 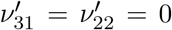. Note that *κ* must be 1 for this example and so the set-point is exactly the ratio *θ*_1_/*θ*_2_. If *m* = 0 this antithetic network is also maxRPA in the stochastic setting (see Theorem 3.5).

**Figure 5:**
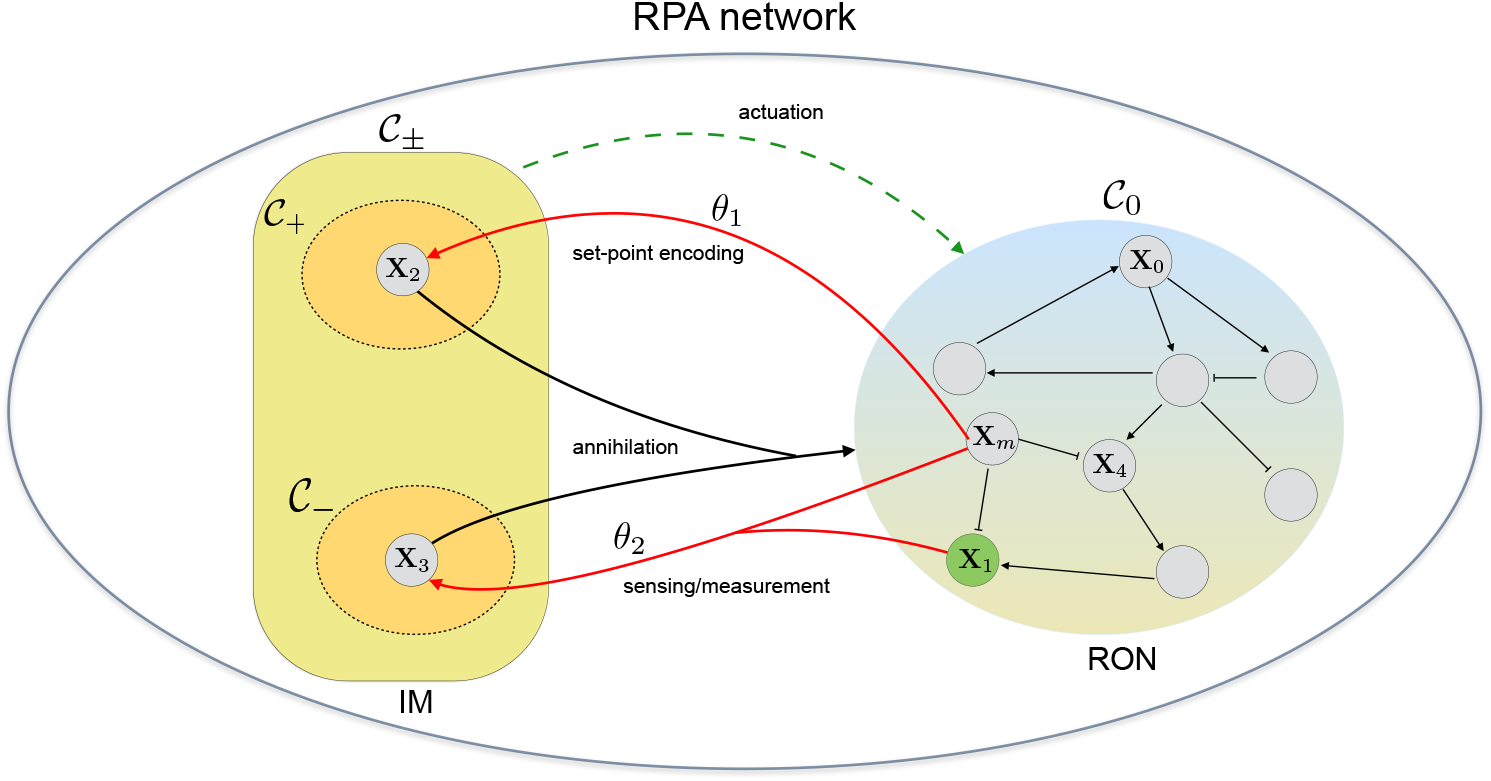
A deterministic antithetic maxRPA network. The internal model (IM) has only two species 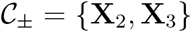 with *q*_2_ = −*q*_3_ > 0. A species **X**_*m*_ (with *m* ≠ 0) belonging to the rest of the network (RON) catalyses the production of these two species via reactions 1 and 2. These two species degrade mutually via a generalised annihilation reaction (4.37). This network is also maxRPA in the stochastic setting if and only if *m* = 0.

**Case 2:** *m* ∈ {2,3}: Without loss of generality we assume that *m* = 2 as the other case (*m* = 3) is similar. Then (3.17) implies that

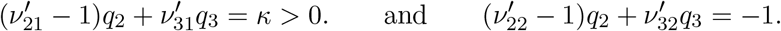

As *q*_2_ > 0 and *q*_3_ < 0, the first reaction must produce at least two molecules of **X**_2_ (i.e. 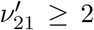). If either 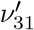 or 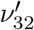 is positive then species **X**_3_ is being produced by these reactions but not being degraded, and so *q*_2_ = −*q*_3_ and the two IM species must degrade mutually via a generalised annihilation reaction (4.37), giving rise to a similar antithetic structure as in the previous case. On the other hand if 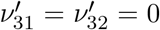, then species **X**_3_ is not getting created by the first two reactions, and we must necessarily have 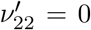, implying that reaction 2 causes degradation of **X**_2_ molecules, without the need of additional annihilation reactions. In this scenario, species **X**_3_ merely acts as a bystander and does not participate in creating the integral action that generates robustness. This integral action is being created only by species **X**_2_ as in the case of homothetic autocatalytic networks with a singleton internal model (see Section 4.1.2).

## 5 Biological examples

The aim of this section is to illustrate our results with known examples of maxRPA networks from existing literature. In the context of synthetic biology, antithetic maxRPA networks can be engineered by interfacing any intracellular network with the antithetic integral feedback (AIF) controller [8]. This was successfully realised in bacteria [24], in vitro [43] and recently in mammalian cells [44]. For *E. Coli*, a slight variant of this approach was adopted in the design of the quasi-integral controller [32] and it is speculated that the AIF controller might be involved in the regulation of housekeeping genes [8]. While it is an ongoing effort to find more examples of endogenous networks that fit the antithetic paradigm, we focus on deterministic homothetic maxRPA networks in the rest of this section. Unless otherwise stated, all reactions follow mass-action kinetics in the upcoming examples.

### 5.1 Bacterial chemotaxis network

Chemotaxis refers to the movement of cells towards chemo-attractants or away from chemo-repellants [4, 30]. In bacterial cells such as *E. Coli* the mechanism of chemotaxis is well-understood and it is characterised by alternating phases of random tumbling and straight-line motion. Upon addition of an attractant the tumbling frequency first decreases sharply but then adapts perfectly to the previous level, demonstrating the phenomenon of RPA. In fact for a simplified model of bacterial chemotaxis (see Figure 6), adapted from [45], this RPA is maximal and we illustrate this using our characterisation result (Theorem 3.2).

**Figure 6:**
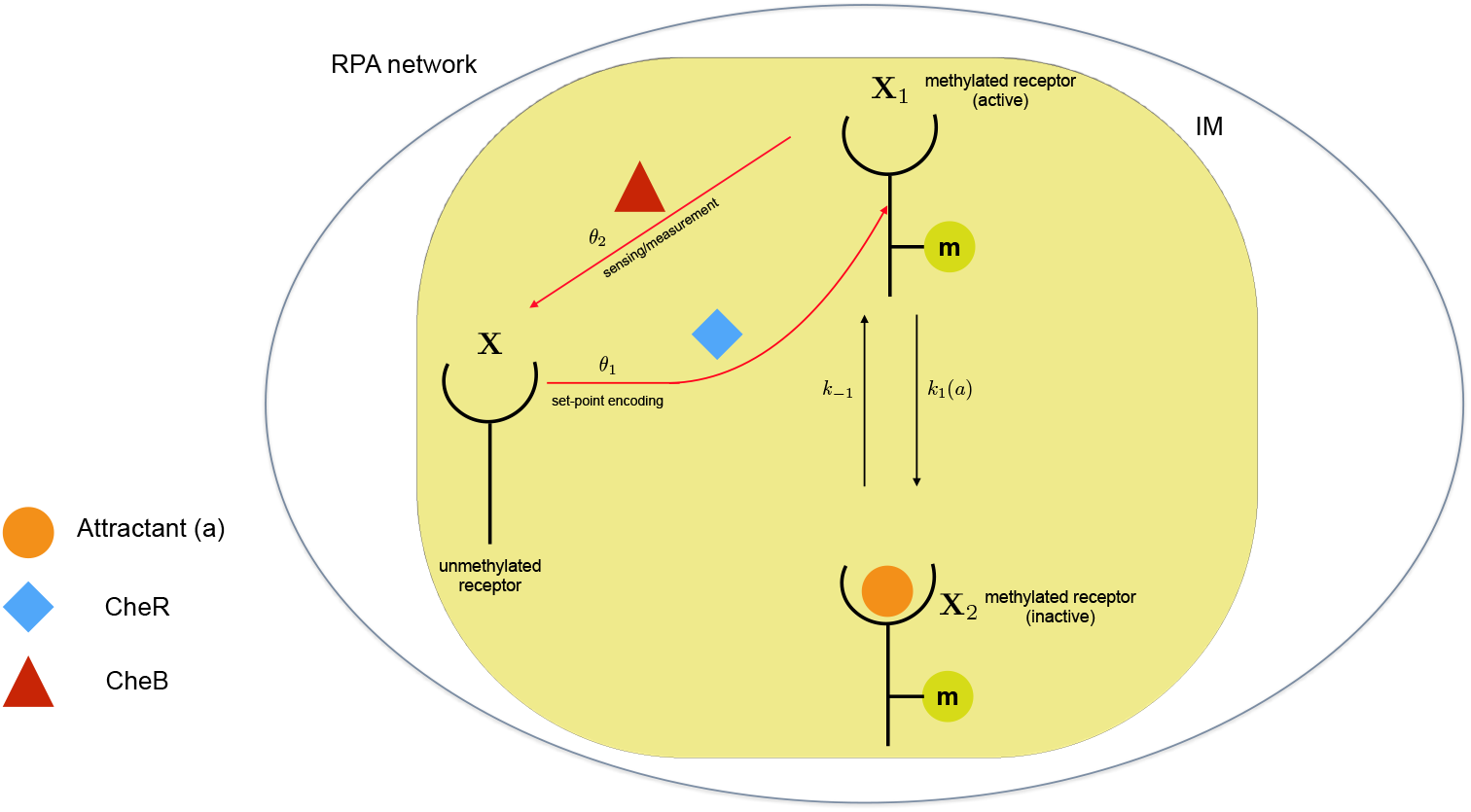
Bacterial chemotaxis network. This figure depicts a simplified model of bacterial chemotaxis adapted from [45]. The two key players are the active and the inactive forms of the methylated receptor, denoted by **X**_1_ and **X**_2_ respectively. In the latter an attractor molecule is bound to the receptor while in the former the receptor is free. The two forms reversibly switch between each other and the rate of switching from **X**_1_ to **X**_2_ depends on the attractor concentration a. In the active form, the methylated receptor **X**_1_ can get demethylated by the phosphatase CheB to produce the deactivated receptor **X**, which in turn can get methylated by enzyme CheR to produce **X**_1_. The RPA output is the concentration of **X**_1_ which determines the bacterial tumbling rate.

The networks consists of three species - the active methylated receptor **X**_1_, the inactive methylated receptor **X**_2_ and the unmethylated receptor **X**. These species interact through four reactions:

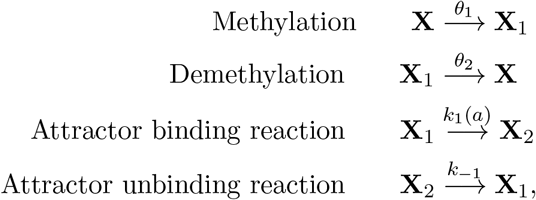

where *θ*_1_ and *θ*_2_ are determined by the concentrations of enzymes CheR and CheB respectively, and *a* denotes the concentration of the attractor. Assuming the unmethylated receptors are in abundance, we can replace **X** by the null species 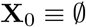 in these reactions to obtain the following 2 × 4 stoichiometry matrix

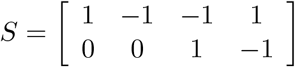

Notice that condition (3.22) holds, and for *q* = (1,1) and *κ* = 1 the linear system (3.17) is also satisfied. Therefore by Theorem 3.2, this network exhibits maxRPA with output set-point *θ*_1_/*θ*_2_ and an integrator is given by the function

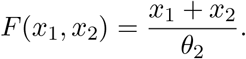

In this example, methylation is the *set-point encoding* reaction, demethylation is the *sensing* or the *measurement* reaction, while the attractor binding and unbinding reactions are the *disturbance inducing* reactions.

Observe that if **X** is not the null species 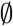, then instead of a zero-order process, the methylation of the unmethylated receptor by CheR becomes a first-order process. In this case, condition (3.22) will get violated and the network will not exhibit maxRPA. This loss of maxRPA also occurs if CheB is allowed to demethylate both active and inactive receptors (**X**_1_ and **X**_2_), as we cannot find a solution to the linear system (3.17) in this case. These observations regarding network constraints for the RPA property have already been made for this chemotaxis network (see [2]), but our characterisation result facilitates drawing such conclusions for arbitrarily complex networks.

### 5.2 Proliferative control of cell lineages

Cell lineages are basic units of tissue and organ development, and the process of lineage proliferation is characterised by several stages of distinct progenitor cells that culminate into mature, highly-differentiated and non-dividing terminal cells that realise various tissue-functions [46]. This process is initiated by pluripotent stem cells and at each stage the progenitor cells can either self-replicate or progress to the next stage, with probabilities depending on the expression of certain stage-specific genes [47]. It is well-known that this lineage proliferation process is tightly-regulated and highly-robust, in the sense that the concentration of terminal cells is insensitive to the stage-specific probabilities that decide if a progenitor cell self-renews or passes to the next stage. This robustness is crucial for development and also for proper tissue repair following an injury.

It was shown computationally and experimentally in [31] that this robustness is achieved when the terminal cells secrete a chemical that can inhibit the self-renewal of a progenitor cell. For maximal robustness, this progenitor cell must be the stem cell and we explore the property of maxRPA by considering an adapted version of the cell-lineage proliferation network given in [31] (see Figure 7). This network consists of *N* species given by **X**_1_,…, **X**_*N*_, with **X**_*N*_ being the stem cell, **X**_1_ being the terminal differentiated cell, and **X**_2_,…, **X**_*N*−1_ being the progenitor cells at (*N* – 2) intermediate stages. These species interact through the following (2*N* – 1) reactions:

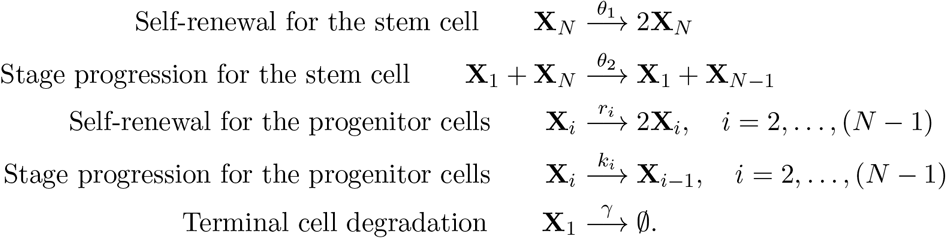

**Figure 7:**
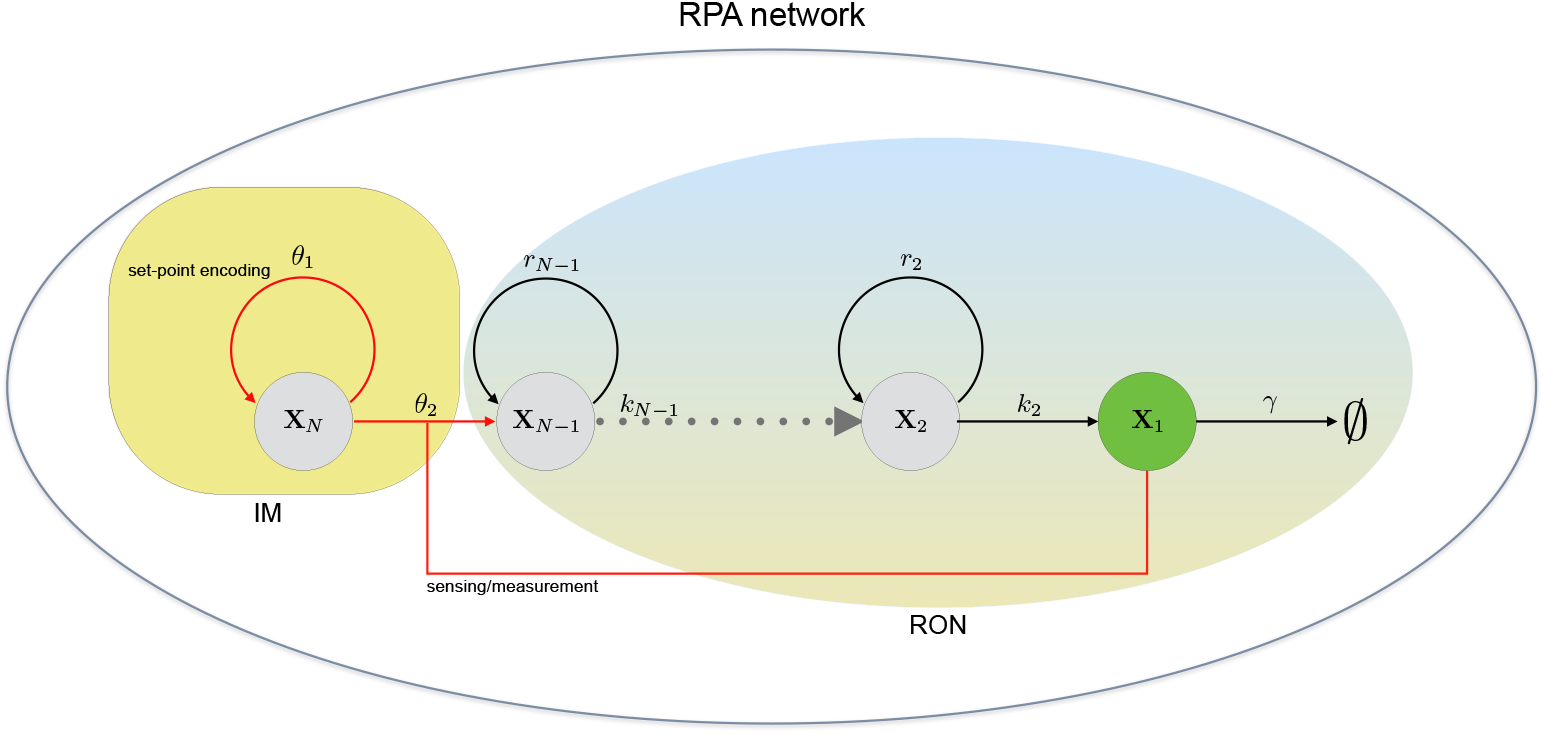
Proliferative network for cell-lineages. This figure depicts a motif adapted from [31] for control of proliferative cell lineages. This *N*-species network begins with a pluripotent stem cell **X**_*N*_, progresses through (*N* – 2) intermediate progenitor stages **X**_*N*−1_,…,**X**_2_ and then ends with a terminal differentiated cell **X**_1_. At each stage (except the last) the cells can *self-replicate* or progress to the next stage. The terminal cell can spontaneously degrade and it can also inhibit the proliferative tendency of the stem cell **X**_*N*_. The RPA output is the concentration of the terminal cell **X**_1_ which determines the size of the tissue in organ development and influences its maintenance and regeneration.

Observe that for any intermediate progenitor cell **X**_*i*_ the chance that it will self-replicate, rather than progress to the next stage, is given by the ratio *r_i_*/(*r_i_* + *k_i_*). For the stem cell this probability becomes

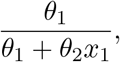

where *x*_1_ is the terminal cell concentration. This clearly shows the negative feedback exerted by the terminal cell on the stem cell by suppressing its tendency to self-replicate.

One can see that condition (3.22) holds and by writing the *N* × (2*N* – 1) stoichiometry matrix one can easily check that for *q* = (1, 0,…, 0) and *κ* = 1 the linear system (3.17) is satisfied. Hence by Theorem 3.2 we can conclude that this network shows maxRPA with output set-point *θ*_1_/*θ*_2_. Therefore the concentration of the terminal cells is robustly maintained at the ratio *θ*_1_/*θ*_2_, and remarkably, it is insensitive to the self-replication and stage-progression rates of the intermediate cells and also the degradation rate of the terminal cell. Observe that this network is an instance of the autocatalytic network consider in Section 4.1.2.

### 5.3 Phosphorylation cycle based integral controller

Our next example is of a synthetic controller that enables an arbitrary intracellular network to achieve maxRPA by implementing the integral feedback mechanism via a phosphorylation cycle [32] (see Figure 8). This cycle constitutes the IM for the maxRPA network with output **X**_1_, while the network being controlled is the RON. The two species in the cycle are the active phosphorylated form of a substrate **X**_2_ and its inactive unphosphorylated forms 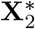. The rates of switching between these two forms obeys Michaelis–Menten enzyme kinetics, as shown in Table 1. Assuming that both forms of the substrate are in high abundance, the reaction 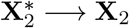 can be viewed as zeroth-order production while the other reaction 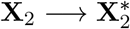 can be viewed as zeroth-order degradation (see Table 1). Under these approximations, this network becomes a zeroth-order degradation network presented in Section 4.1.1. The IM consists of a single species **X**_2_ and as per Theorem 3.2, the network exhibits maxRPA for the output species **X**_1_ which represents the phosphatase enzyme that is responsible for catalysing the dephosphorylation reaction.

**Figure 8:**
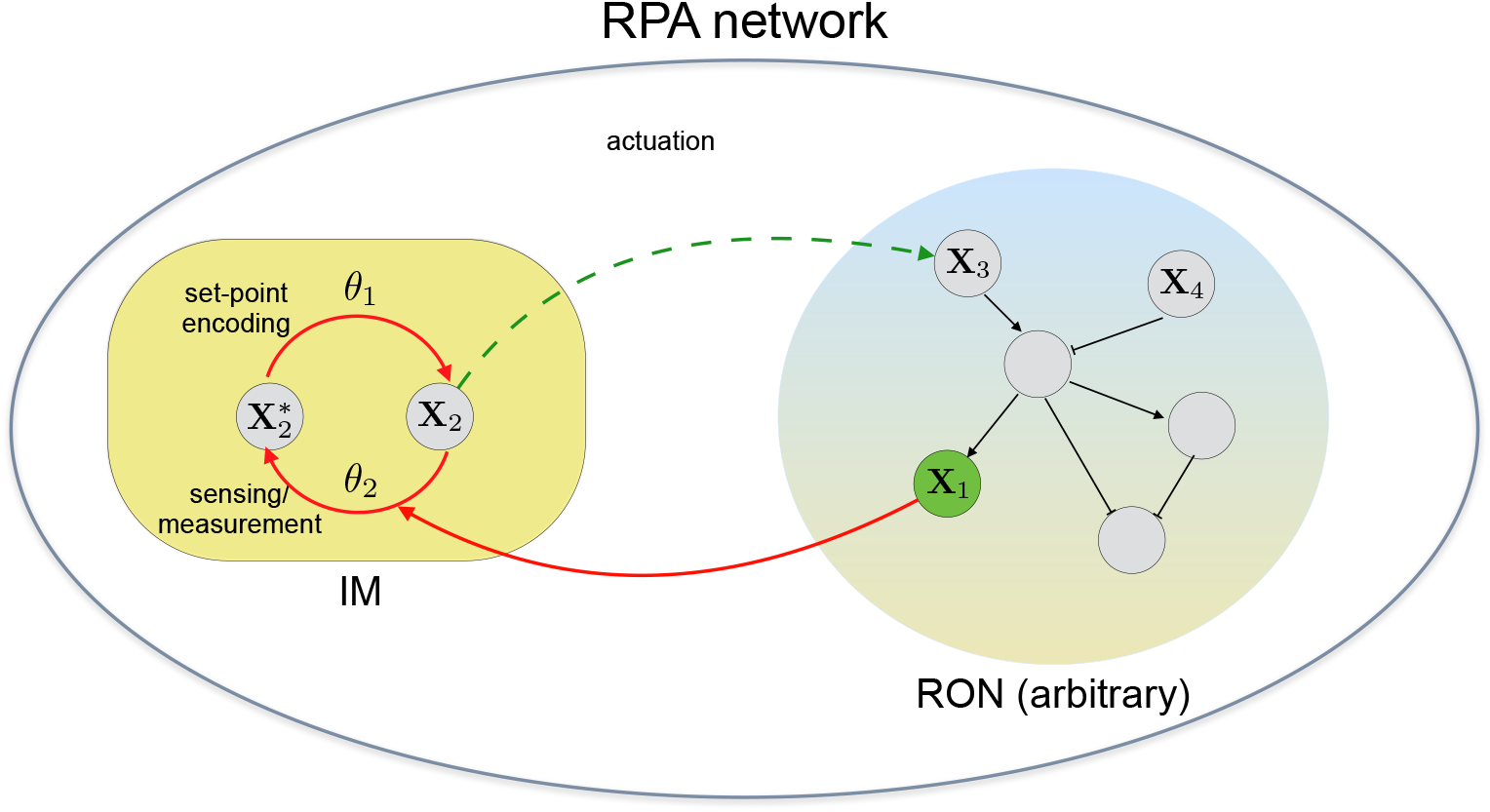
Integral controller based on phosphorylation cycle. This figure depicts the scenario, adapted from [32], where an arbitrary network (RON) is controlled by a module (IM) comprising a substrate in phosphorylated **X**_2_ and unphosphorylated 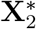 forms. The rate constant *θ*_1_ for the phosphorylation reaction is determined by the kinase concentration, while the dephosphorylation reaction occurs at a rate that is proportional to the concentration of the phosphatase **X**_1_ which is the RPA output. The arbitrary network (RON) gets actuated by the phosphorylated substrate **X**_2_ and it produces the phosphatase **X**_1_.

**Table 1:** Reactions in the Phosphorylation cycle. Both reactions follow Michaelis–Menten kinetics with Michaelis constants *K*_1_ and *K*_2_, and *x*_1_, *x*_2_ and 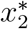 denote the concentrations of **X**_1_, **X**_2_ and 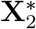 respectively. For the phosphorylation reaction, the kinase concentration is fixed and *θ*_1_ is its product with the catalytic rate constant of this enzyme. For the dephosphorylation reaction, *θ*_1_ is the catalytic rate constant of the phosphatase. When *x_2_* and 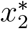 have high values the two reactions and their propensities can be approximated as indicated in the last two columns.

| No. | Name | Reaction | Propensity | Approx.<br>Reaction | Approx.<br>Propensity |
| --- | --- | --- | --- | --- | --- |
| 1 | Phosphorylation | $\mathbf{X}_2^* \longrightarrow \mathbf{X}_2$ | $\frac{\theta_1 x_2^*}{x_2^* + K_1}$ | $\emptyset \longrightarrow \mathbf{X}_2$ | $\theta_1$ |
| 2 | Dephosphorylation | $\mathbf{X}_2 \longrightarrow \mathbf{X}_2^*$ | $\frac{\theta_2 x_1 x_2}{x_2 + K_2}$ | $\mathbf{X}_2 \longrightarrow \emptyset$ | $\theta_2 x_1$ |

### 5.4 The sniffer system

The *sniffer system* [33] (see Figure 9) exemplifies an important regulatory motif called the *incoherent feedforward loop* (IFFL). This motif is found in many biological networks [45] and particularly in signalling networks [48, 49]. In an IFFL the input node influences the output node via two paths that have the opposite effect, i.e. one path activates the production of the output while the other represses it. The structure of IFFL ensures that the cumulative influence of the input node on the output node *cancels out*, thereby ensuring RPA w.r.t all disturbances that only affect the input node.

**Figure 9:**
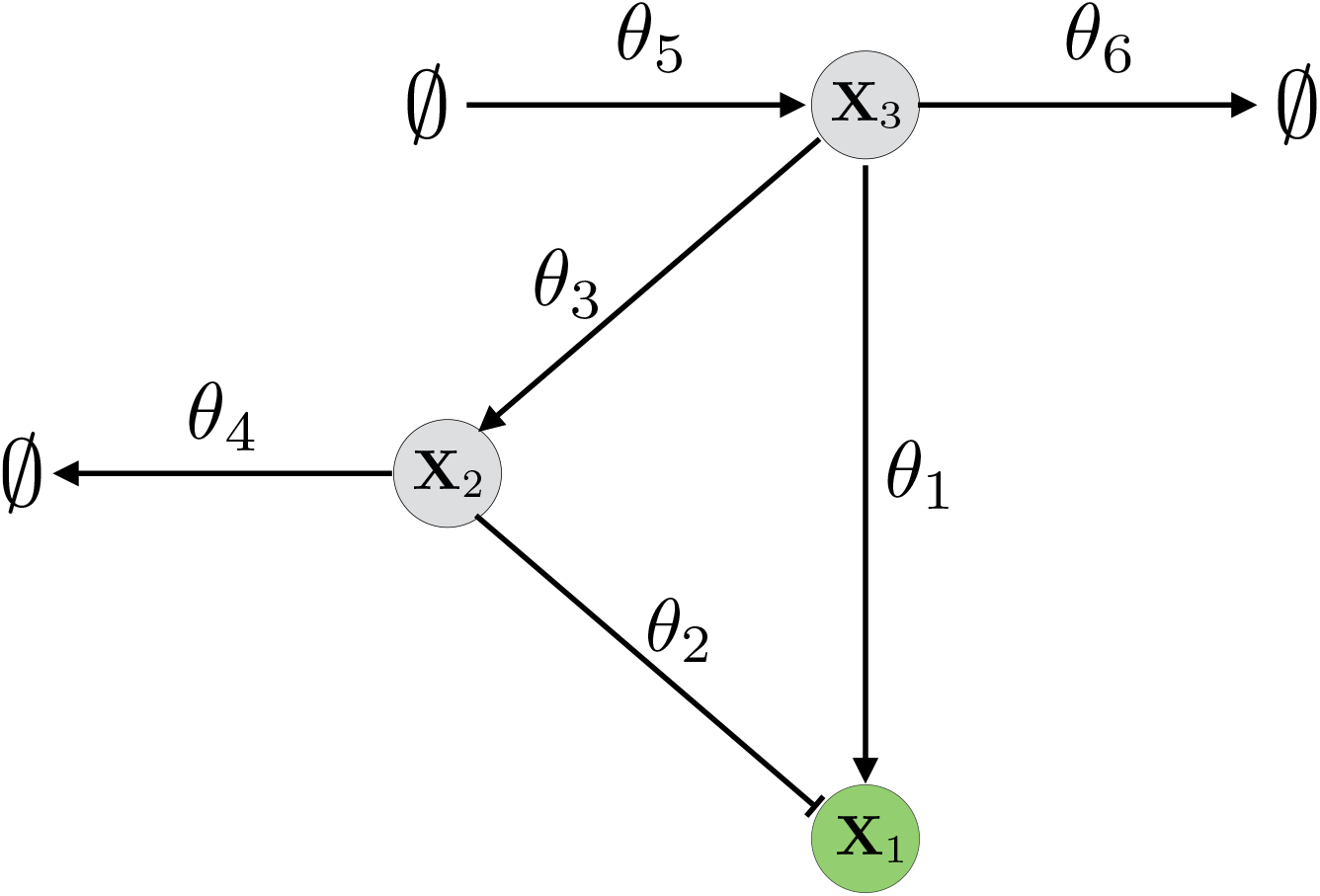
Sniffer system. This figure depicts the sniffer system, adapted from [33], which comprises an *incoherent feedforward loop* (IFFL) from the input **X**_3_ to the output **X**_1_. The input directly produces the output, but it also produces another species **X**_2_ that represses the output by causing its degradation. The reactions of the network are given by (5.40). All the reactions follow mass-action kinetics with rate constants as indicated in the figure.

In the sniffer system shown in Figure 9, the input node is given by species **X**_3_, species **X**_1_ is the output node, while another species **X**_2_ serves as an intermediate node. The input species **X**_3_ catalytically produces both **X**_1_ and **X**_2_, and **X**_2_ catalytically degrades **X**_1_. We include inflow and outflow reactions for **X**_3_ and also an outflow reaction for **X**_2_. Overall this three-species network has the following six reactions:

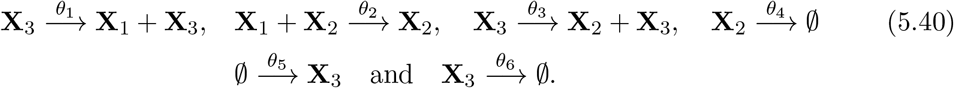

The stoichiometry matrix for this network is

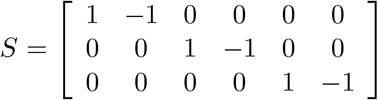

and it can be seen that for *q* = (1, 0, 0) and *κ* = 1, condition (3.17) holds. However the other stoichiometric condition stipulated by Theorem 3.2 does not hold and hence this network will not exhibit maxRPA. This can also be verified independently by noting that the steady-state values of the species concentrations are given by

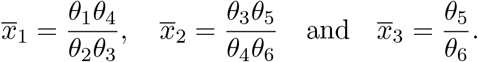

Hence the output level 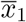 is only robust to perturbations in parameters *θ*_5_ and *θ*_6_ that directly affect the input, while it is sensitive to all the other parameters that determine the flow of information from the input to the output.

## 6 Conclusion

Intracellular biomolecular networks are immensely complex and yet they must tightly regulate key molecular players — failing to do so can have catastrophic ramifications for the cell-population and consequently for the whole organism [3]. As these networks operate in an unpredictable and fluctuating environment, it is conceivable that they have evolved to attain a form of “maximal robustness”, whereby the output being regulated is sensitive to the least number of networks parameters, and insensitive to maximum number of disturbance sources which can be both parametric or structural. The goal of this paper is to develop a mathematical theory for networks exhibiting *maximal robust perfect adaptation* (maxRPA). We consider both deterministic and stochastic descriptions of the dynamics and provide necessary and sufficient conditions for maxRPA. These conditions pertain to only the stoichiometric structure of the network and they are linear-algebraic in nature, and hence they can be easily checked for large dimensional networks.

Our results are best understood from the control-theoretic viewpoint offered by the well-known *internal model principle* (IMP) (see [28]). It stipulates that for a network to achieve robustness, a subnetwork, called the *internal model* (IM), must measure the deviation of the output from its set-point, and then pass its time-integral as an actuating signal to the *rest of the network* (RON) in a feedback fashion (see Figure 1). We formalise this principle in the context of maxRPA networks and demonstrate how the IM arises naturally from our stoichiometric conditions that characterise such networks. One can interpret the IM as an embedded controller module that ensures robustness, thereby connecting our results with the synthetic biology paradigm for designing controller modules for generating RPA [8, 24, 32]. An important question is to determine the minimal complexity of the IM that is needed for maxRPA. Using our characterisation results we answer this question fully in both deterministic and stochastic settings and we highlight the differences between the two scenarios. We also illustrate our results with known examples of maxRPA networks in the existing literature.

Existing RPA studies typically assume that disturbances can enter the dynamics via a dedicated input node [18, 19], and in such a setting only two possible RPA network topologies can arise - namely the *incoherent feedforward* (IFF) topology and the *negative feedback* (NFB) topology. In contrast, in this study of maxRPA networks, disturbances can enter from multiple nodes and they can vary the parameters and also alter the network structure. Therefore the set of allowed disturbances is significantly larger, and consequently, maxRPA networks cannot have the IFF topology as here robustness is engendered via a balancing mechanism between disparate output affecting network branches — and this balancing mechanism can be disrupted by the disturbances leading to the loss of RPA (see the example in Section 5.4). As our results show, all maxRPA networks must have the NFB topology, and in addition, the stoichiometric structure of the network must satisfy a certain linear-algebraic condition.

We believe that the notion of maxRPA is important for understanding and designing robust biomolecular networks, especially if robustness needs to be preserved within the cellular milieu which is rife with disturbances. These disturbances include fluctuations in extrinsic factors, such as nutrient concentration, temperature, pressure etc. [1], or intrinsic factors, such as, metabolic burden [21, 22], cross-talk with other cellular components [20], loading effects [23, 40]. In synthetic biology applications, our results can facilitate modular design by providing a framework for building orthogonal components that behave predictably upon connection due to robustness against the connection-induced disturbances [50]. The theory we have developed is in the idealised scenario where the adaptation is perfect and all disturbances that do not affect the output set-point are rejected. However, our results provide a starting point for finding or constructing realistic biological networks that are *nearly* maxRPA. As illustrated by the example in Section 5.3, such networks would *approximately* satisfy the required stoichiometric conditions, under some reasonable assumptions on the abundance levels of the constituent species.

## Acknowledgments

This project has received funding from the European Research Council (ERC) under the European Union’s Horizon 2020 research and innovation programme grant agreement no. 743269 (CyberGenetics project).

## Footnotes

1 The irreducibility condition demands that for any two states *x*, 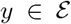 there must exist a positiveprobability sequence of reactions *k*_1_,…, *k_n_* that takes the state *x* to the state *y*.

2 Such a function demonstrates that the dynamics has an attractive tendency towards a compact subset of the state-space.

