## Supplementary Text for "Universal structural requirements for maximal robust perfect adaptation in biomolecular networks"

**S1 Stochastic Reaction Networks** **3**

**S2 Encoding of the set-point** **5**

**S3 Proofs of the Main Results** **5**

### Supplementary Text

#### S1 Stochastic Reaction Networks

In this section we supplement Section 2 in the main text with some additional information on stochastic reaction networks. Recall that in the *continuous-time Markov chain* (CTMC) model of a reaction network [1], the state $x = (x_1, \dots, x_N) \in \mathbb{N}_0^N$ is the vector of species copy-numbers or molecular counts, the propensity function $\lambda_k : \mathbb{N}_0^N \to \mathbb{R}_+$ maps the state $x$ to the rate of firing $\lambda_k(x)$ of reaction $k$, and $\zeta_k \in \mathbb{Z}^N$ denotes the state-displacement caused by reaction $k$. The CTMC $(X(t))_{t\geq 0}$ for the reaction dynamics can be characterised by its *generator* [2] defined as

$$\mathbb{A}f(x) = \sum_{k=1}^{K} \lambda_k(x)(f(x+\zeta_k) - f(x)), \tag{S1}$$

where $f$ is a bounded real-valued function on the set $\mathcal{E} \subset \mathbb{N}_0^N$ of accessible states for the CTMC. Under certain special conditions, the domain of the generator can be taken to be the set of all polynomially growing functions on the state-space $\mathcal{E}$ [3]. Letting $p_t$ to be probability distribution of $X(t)$ i.e.

$$p_t(x) = \mathbb{P}(X(t) = x), \quad x \in \mathcal{E},$$

the well-known *Chemical Master Equation* (CME) [1] expresses the time-evolution of $p_t$ as

$$\frac{dp_t}{dt} = \mathbb{A}^* p_t, \tag{S2}$$

where $\mathbb{A}^*$ is the *adjoint* of linear operator $\mathbb{A}$ given by

$$\mathbb{A}^*\xi(x) = \sum_{k=1}^{K} \left(\lambda_k(x-\zeta_k)\xi(x-\zeta_k) - \lambda_k(x)\xi(x)\right).$$

The CTMC $(X(t))_{t\geq 0}$ is said to be *ergodic* if regardless of the initial distribution, as $t \to \infty$, the probability distribution $p_t$ converges to a stationary distribution $\pi$ in the $\ell_1$ norm i.e.

$$\lim_{t\to\infty} \|p_t - \pi\|_{\ell_1} := \lim_{t\to\infty} \sum_{x\in\mathcal{E}} |p_t(x) - \pi(x)| = 0.$$

If this convergence is exponential i.e. there exists a constant $\rho > 0$ and another constant $C > 0$ (depending on $p_0$), such that for any $t > 0$

$$\|p(t) - \pi\|_{\ell_1} \leq Ce^{-\rho t}, \tag{S3}$$

then the CTMC is called *exponentially ergodic* [4]. The stationary distribution $\pi$ is a globally attracting fixed point for the CME (S2), over the space of probability distributions over $\mathcal{E}$, and it satisfies

$$\mathbb{A}^*\pi = \mathbf{0}. \tag{S4}$$

In order to verify ergodicity, a two-step approach is needed. First one needs to check if the state-space $\mathcal{E}$, which is typically infinite, is *irreducible* and then construct a suitable Foster-Lyapunov function over the state-space $\mathcal{E}$ [4]. In order for the state-space $\mathcal{E}$ to be irreducible, for any two states $x, y \in \mathcal{E}$ there must exist a positive-probability sequence of reactions $k_1, \dots, k_n$ that take state $x$ to $y$. In light of condition (2.5) this is equivalent to having

$$z_{j-1} \geq \nu_{k_j} \qquad \text{for each} \qquad j = 1, \dots, n$$

where $z_0 = x$, $z_n = y$ and

$$z_j = z_0 + \sum_{l=1}^{j} \zeta_{k_l} \qquad \text{for} \qquad j = 1, \dots, n.$$

A Foster-Lyapunov function $V : \mathcal{E} \to [1, \infty)$ on the state-space $\mathcal{E}$ essentially shows that the CTMC has an attractive tendency toward a compact set [4]. To verify exponential ergodicity this function needs to be *norm-like* (i.e. all sub-level sets must be compact) and for some $C_1, C_2 > 0$

$$\mathbb{A}V(x) \leq C_1 - C_2 V(x) \qquad \text{for all} \qquad x \in \mathcal{E}, \tag{S5}$$

where $\mathbb{A}$ is the generator of the CTMC (see Theorem 7.1 in [4]). For stochastic reaction networks, computational frameworks for systematically checking state-space irreducibility and constructing Foster-Lyapunov functions are provided in [5] and [3]. In fact, for many biological networks, a linear Foster-Lyapunov

$$V(x) = 1 + \langle v, x \rangle \tag{S6}$$

can be constructed, where $\langle \cdot, \cdot \rangle$ denotes the standard inner product on $\mathbb{R}^N$ and $v \in \mathbb{R}^N_+$ is a vector found by linear programming [3].

For any real-valued function $f(x)$ over the state-space $\mathcal{E}$ satisfying

$$\|f\|_V := \sup_{x \in \mathcal{E}} \frac{|f(x)|}{V(x)} < \infty \tag{S7}$$

we have the ergodic convergence (see Theorem 6.1 in [4])

$$\lim_{t \to \infty} \mathbb{E}(f(X(t))) = \mathbb{E}_\pi(f) := \sum_{x \in \mathcal{E}} f(x)\pi(x), \tag{S8}$$

where $\mathbb{E}$ denotes the expectation operator and $\mathbb{E}_\pi$ denotes expectation under the stationary distribution. Moreover (S4) implies that

$$\mathbb{E}_\pi(\mathbb{A}f) = 0. \tag{S9}$$

### S2 Encoding of the set-point

Recall from Section 2.2 that we suppose that the first $m$ reactions have mass-action kinetics and their rate constants (denoted by $\theta_1, \dots, \theta_m$) encode the set-point $\theta^*$ via some smooth function $\phi$, i.e.

$$\phi(\theta_1, \dots, \theta_m) = \theta^*.$$

If we multiply all the network propensity functions by a positive scalar $r$, then the steady-state will not change (in both deterministic and stochastic settings) as we are essentially just recalibrating time. Hence

$$\phi(r\theta_1, \dots, r\theta_m) = \phi(\theta_1, \dots, \theta_m).$$

Differentiating this relation w.r.t. $r$ we obtain the following first-order *partial differential equation* (PDE)

$$\theta_1 \frac{\partial \phi}{\partial \theta_1} + \cdots + \theta_m \frac{\partial \phi}{\partial \theta_m} = 0.$$

Note that if $m = 1$ then the only possible solution is that $\phi$ is a constant function. This is not feasible as this would imply that the set-point does not depend on any of the propensity functions! Hence we look for a solution with $m = 2$ of the PDE

$$\theta_1 \frac{\partial \phi}{\partial \theta_1} + \theta_2 \frac{\partial \phi}{\partial \theta_2} = 0.$$

The *characteristic curves* for this PDE, with initial values $(\theta_1, \theta_2)$ are $\theta_1(s) = \theta_1 e^s$ and $\theta_2(s) = \theta_2 e^s$. Along these curves the function $\phi$ is constant, i.e.

$$\phi(\theta_1 e^s, \theta_2 e^s) = \phi(\theta_1, \theta_2).$$

Notice that this also proves that for any $\theta_1, \theta_2 > 0$

$$\phi(\theta_1, \theta_2) = \phi\left(\frac{\theta_1}{\theta_2}, 1\right).$$

Hence the set-point encoding function $\phi$ is essentially a function of only one variable, which is the ratio of $\theta_1$ and $\theta_2$. This function is denoted by $\phi_{\text{out}}$ in the main text.

### S3 Proofs of the Main Results

In this section we present the proofs of our main results in the paper. These results provide the necessary and sufficient linear-algebraic conditions for a system to achieve maxRPA, in both deterministic and stochastic settings. Recall that $S$ is the $N \times K$ stoichiometric matrix for the reaction network given by (3.15) and we shall assume that it has full row-rank (3.16). Next we state and prove a couple of simple lemmas.

**Lemma S3.1** *Consider a network satisfying our stability assumption (i.e. Assumption 3.1(A) in the deterministic setting or Assumption 3.4(A) in the stochastic setting). Let $S$ be its stoichiometric matrix satisfying* (3.16)*. Then there can exist at most one solution $(q,\kappa)$ for which* (3.17) *is satisfied and this solution can be found by solving a linear system of the form $Ax = b$.*

**Proof.** Let us first assume that we are in the deterministic setting and fix a nominal value of $(\theta^*, \lambda^*) \in \Theta \times \Lambda_u$ for which the steady-state is $\overline{x} = \overline{x}_{\theta^*,\lambda^*}$. Let $c = (c_1, \dots, c_K)$ be the vector of all the propensity functions at this steady-state, i.e.

$$c = (\theta_1^* m_1(\overline{x}), \theta_2^* m_2(\overline{x}), \lambda_3^*(\overline{x}), \dots, \lambda_K^*(\overline{x})).$$

The fixed-point relation (2.2) implies that

$$Sc = \mathbf{0}. \tag{S10}$$

Since the fixed-point $\overline{x}$ is in the positive orthant and $m_1(x)$ and $m_2(x)$ are mass-action monomials we must have that both $c_1$ and $c_2$ are strictly positive. Now suppose that there are two pairs $(q_1, \kappa_1)$ and $(q_2, \kappa_2)$ satisfying (3.17). Then we have

$$0 = q_i^T Sc = (\kappa_i, -1, 0, \dots, 0)c = \kappa_i c_1 - c_2 \quad \text{for} \quad i = 1, 2.$$

Therefore $\kappa_1 = \kappa_2 = c_2/c_1$ and since the matrix $S$ has full row-rank we must also have $q_1 = q_2$. This proves the uniqueness of the solution of the linear system (3.17) if it exists. Checking this existence is equivalent to checking the existence of a solution to the linear system $Ax = b$ where $x = (q, \kappa)$ is the $(N+1)$ dimensional column vector of unknowns, $b$ is the $K$ dimensional vector given by $b = (0, -1, 0, \dots, 0)$ and $A$ is the $K \times (N+1)$ dimensional matrix obtained by appending $-e_1$ as a column to the transpose of $S$. Here $e_1$ denotes the $N$ dimensional vector with the first component as one and the rest of the components as zeros.

The proof in the stochastic case is similar except that $c$ is the vector of expected propensity functions of all the reactions at the stationary distribution. Relation (S10) can be obtained from (S9) by setting $f(x) = x$, i.e. the identity function of the state vector. □

**Lemma S3.2** *Consider a maxRPA network characterised by the solution $(q, \kappa)$ to the linear-algebraic system* (3.17)*. If* (3.31) *holds then $\mathbf{X}_1 \notin \mathcal{C}_\pm$ and hence the output species $\mathbf{X}_1$ belongs to $\mathcal{C}_0$ and it is not present in the internal model.*

**Proof.** Condition (3.31) implies the existence of a vector $c = (c_1, \dots, c_K)$ such that $c_1 = c_2 = 0$ and

$$Sc = e_1$$

where $S$ is the stoichiometry matrix for the network. Multiplying (3.17) by the $K \times 1$ vector $c$ on the right we see that

$$0 = (\kappa, -1, 0, \dots, 0)c = q^T Sc = q^T e_1 = q_1.$$

Hence $\mathbf{X}_1 \notin \mathcal{C}_0$ and this completes the proof of this lemma. □

#### S3.1 Characterisation of deterministic maxRPA networks

**Proof.**[Proof of Theorem 3.2] Fix a $(\theta, \lambda) \in \Theta \times \Lambda_u$ and for any $\gamma = (\gamma_1, \dots, \gamma_R) \in (-\epsilon, \epsilon)^R$ let

$$\lambda_\gamma = \lambda + \sum_{i=1}^{R} \gamma_i \phi_i$$

where $\phi_1, \dots, \phi_R$ are as in Assumption 3.1(B). Due to Assumption 3.1(A) there is a fixed point $\overline{x}_{\theta,\lambda_\gamma}$ satisfying

$$D_{\theta,\lambda_\gamma}(\overline{x}_{\theta,\lambda_\gamma}) = \mathbf{0}. \tag{S11}$$

Differentiating this jointly w.r.t. parameters $(\theta, \gamma)$ gives us

$$\mathcal{J}_x D_{\theta,\lambda_\gamma}(\overline{x}_{\theta,\lambda_\gamma}) \mathcal{J}_{\theta,\gamma} \overline{x}_{\theta,\lambda_\gamma} + \mathcal{J}_{\theta,\gamma} D_{\theta,\lambda_\gamma}(\overline{x}_{\theta,\lambda_\gamma}) = \mathbf{0} \tag{S12}$$

where $\mathcal{J}_\alpha$ denotes the Jacobian w.r.t. $\alpha$. Note that $\lambda_{\mathbf{0}} = \lambda$, the fixed point $\overline{x}_{\theta,\lambda}$ is globally attracting, and the matrix $\mathcal{J}_x D_{\theta,\lambda}(\overline{x}_{\theta,\lambda})$ is invertible. Setting $\gamma = \mathbf{0}$ and multiplying relation (S12) by the inverse of this Jacobian matrix on the left we obtain

$$\mathcal{J}_{\theta,\gamma} \overline{x}_{\theta,\lambda_\gamma} \Big|_{\gamma=\mathbf{0}} = - \left[ \mathcal{J}_x D_{\theta,\lambda}(\overline{x}_{\theta,\lambda}) \right]^{-1} \mathcal{J}_{\theta,\gamma} D_{\theta,\lambda_\gamma}(\overline{x}_{\theta,\lambda}) \Big|_{\gamma=\mathbf{0}}. \tag{S13}$$

From the form of the vector field $D_{\theta,\lambda_\gamma}(x)$ (see (3.19)) it is evident that we can write

$$\mathcal{J}_{\theta,\gamma} D_{\theta,\lambda_\gamma}(\overline{x}_{\theta,\lambda}) \Big|_{\gamma=\mathbf{0}} = S \Psi(\overline{x}_{\theta,\lambda}), \tag{S14}$$

where $S$ is the stoichiometry matrix (3.15) and $\Psi(x)$ is the $K \times (R+2)$ dimensional matrix given by

$$\Psi(x) = \begin{bmatrix} m_1(x) & 0 & \mathbf{0} \\ 0 & m_2(x) & \mathbf{0} \\ \mathbf{0} & \mathbf{0} & \Phi(x) \end{bmatrix},$$

where $\Phi(x)$ is the matrix given by (3.21).

Let $f$ be the projection map (2.9) that maps the state vector to the state of the output species. For (2.10) to hold we must have

$$[\nabla_{\theta,\gamma} f(\overline{x}_{\theta,\lambda_\gamma})]^T = \phi'_{\text{out}} \left( \frac{\theta_1}{\theta_2} \right) \left( \frac{1}{\theta_2}, -\frac{\theta_1}{\theta_2^2}, \mathbf{0} \right). \tag{S15}$$

Let $e_1$ denote the vector in $\mathbb{R}^N$ with the first component as 1 and the rest as zeros. From (S13) and (S14), at $\gamma = \mathbf{0}$ we obtain

$$[\nabla_{\theta,\gamma} f(\overline{x}_{\theta,\lambda_\gamma})]^T = e_1^T \mathcal{J}_{\theta,\gamma} \overline{x}_{\theta,\lambda_\gamma} \Big|_{\gamma=\mathbf{0}} = -e_1^T \left[ \mathcal{J}_x D_{\theta,\lambda}(\overline{x}_{\theta,\lambda}) \right]^{-1} S \Psi(\overline{x}_{\theta,\lambda}). \tag{S16}$$

Letting

$$q_{\theta,\lambda}^T = -e_1^T \left[\mathcal{J}_x D_{\theta,\lambda}(\overline{x}_{\theta,\lambda})\right]^{-1} \tag{S17}$$

and equating (S15) and (S16) we get

$$q_{\theta,\lambda}^T \zeta_1 = \frac{\phi'_{\text{out}}\left(\frac{\theta_1}{\theta_2}\right)}{\theta_2 m_1(\overline{x}_{\theta,\lambda})}, \quad q_{\theta,\lambda}^T \zeta_2 = -\frac{\theta_1 \phi'_{\text{out}}\left(\frac{\theta_1}{\theta_2}\right)}{\theta_2^2 m_2(\overline{x}_{\theta,\lambda})} \tag{S18}$$

$$\text{and} \quad q_{\theta,\lambda}^T \widehat{S}\Phi(\overline{x}_{\theta,\lambda}) = \mathbf{0}, \tag{S19}$$

where $\widehat{S} = \text{Col}(\zeta_3, \ldots, \zeta_K)$ contains the last $(K-2)$ columns of the stoichiometry matrix $S$. Due to Assumption 3.1(B), matrix $\Phi(\overline{x}_{\theta,\lambda})$ has full row rank and therefore (S19) can be simplified to

$$q_{\theta,\lambda}^T \zeta_k = 0 \qquad \text{for all} \qquad k = 3, \ldots, K. \tag{S20}$$

Let $c$ be a vector satisfying (S10). Using (S18) and (S20) we obtain

$$0 = q_{\theta,\lambda}^T S c = \phi'_{\text{out}}\left(\frac{\theta_1}{\theta_2}\right)\left[\frac{c_1}{\theta_2 m_1(\overline{x}_{\theta,\lambda})} - \frac{\theta_1 c_2}{\theta_2^2 m_2(\overline{x}_{\theta,\lambda})}\right],$$

and since $\phi'_{\text{out}}\left(\frac{\theta_1}{\theta_2}\right) \neq 0$ (see (2.8)), this shows that

$$\frac{c_1}{c_2} = \frac{\theta_1 m_1(\overline{x}_{\theta,\lambda})}{\theta_2 m_2(\overline{x}_{\theta,\lambda})} = \phi_{\text{out}}^{-1}(\overline{x}_{\theta,\lambda,1}) \prod_{i=1}^{N} \overline{x}_{\theta,\lambda,i}^{(\nu_{i1}-\nu_{i2})} \tag{S21}$$

where the last equality follows from (2.10). Here $\phi_{\text{out}}^{-1}(\cdot)$ denotes the inverse of the set-point encoding function and $\overline{x}_{\theta,\lambda,i}$ is the $i$-th component of the steady-state vector $\overline{x}_{\theta,\lambda}$. Observe that (S21) must hold for all $(\theta,\lambda) \in \Theta \times \Lambda_u$ with the same constant $c_1/c_2$.

From (S13) and (S14) we note that

$$J_{\theta,\lambda} := \mathcal{J}_{\theta,\gamma}\overline{x}_{\theta,\lambda_\gamma}\Big|_{\gamma=\mathbf{0}} = -\left[\mathcal{J}_x D_{\theta,\lambda}(\overline{x}_{\theta,\lambda})\right]^{-1} S\Psi(\overline{x}_{\theta,\lambda}).$$

Due to (3.16) and Assumption 3.1(B), matrices $S$ and $\Psi(\overline{x}_{\theta,\lambda})$ have full row-ranks, and since this property does not change upon multiplication by a nonsingular matrix on the left, we can conclude that matrix $J_{\theta,\lambda}$ also has full row-rank

$$\text{Rank}(J_{\theta,\lambda}) = N. \tag{S22}$$

Applying the *constant rank theorem* (see Theorem 11.1 and Proposition 11.7 in [6]) we can conclude that the map $(\theta,\gamma) \mapsto \overline{x}_{\theta,\lambda_\gamma}$ is locally open and surjective, i.e. it maps an open set $O \subset \Theta \times (-\epsilon,\epsilon)^R$ around $(\theta,\mathbf{0})$ to an open set $P \subset \mathcal{E}$ around $\overline{x}_{\theta,\lambda}$. As relation (S21) holds for each $x \in P$, we must have that $\nu_{i2} = \nu_{i1}$ for each $i = 2, \ldots, N$ and

$$\phi_{\text{out}}^{-1}(x) = \kappa^{-1} x^{\alpha}$$

for $\alpha = \nu_{12} - \nu_{11}$ and $\kappa = \frac{c_2}{c_1} > 0$. Setting

$$q = \frac{\theta_2 m_1(\overline{x}_{\theta,\lambda}) q_{\theta,\lambda}}{\phi'_{\text{out}}\left(\frac{\theta_1}{\theta_2}\right)} \tag{S23}$$

we see that the pair $(q, \kappa)$ must satisfy the linear system (3.17). As no other such pair can exist under our assumptions (see Lemma S3.1) $(q, \kappa)$ does not depend on the choice of $(\theta, \lambda) \in \Theta \times \Lambda_u$. This proves the "only if" part of the result.

The "if" part of the result is trivial as the stability of the deterministic dynamics along with the presence of an integrator (3.24) (see also Remark 3.3) shows that the network exhibits maxRPA. This completes the proof of this theorem. □

#### S3.2 Characterisation of stochastic maxRPA networks

In the stochastic setting, the mass-action propensities for the first two reactions, with rate-constants $\theta = (\theta_1, \theta_2) \in \Theta$, can be written as $\lambda_k(x) = \theta_k m_k(x)$ for $k = 1, 2$, where $m_k$ denotes the combinatorial factor

$$m_k(x) = \prod_{i=1}^{N} \frac{x_i(x_i - 1)\dots(x_i - \nu_{ki} + 1)}{\nu_{ki}!}. \tag{S24}$$

As in the deterministic scenario, the system is affected by disturbances that perturb the propensity map $\lambda$ which belongs to the uncertain set $\Lambda_u$. The stochastic reaction dynamics is represented by a CTMC $(X_{\theta,\lambda}(t))_{t\geq 0}$ with state-space $\mathcal{E} \subset \mathbb{N}_0^N$ and generator $\mathbb{A}_{\theta,\lambda}$ given by

$$\mathbb{A}_{\theta,\lambda} f(x) = \sum_{k=1}^{2} \theta_k m_k(x) \Delta_k f(x) + \sum_{k=3}^{K} \lambda_k(x) \Delta_k f(x), \tag{S25}$$

where $f$ is a real-valued function on the state-space $\mathcal{E}$ and $\Delta_k$ is the difference operator

$$\Delta_k f(x) = f(x + \zeta_k) - f(x).$$

As discussed in Section 2.3, for maxRPA to hold in the stochastic setting, there exists a real-valued function $F_{\theta,\lambda}$ on the state-space $\mathcal{E} = \mathbb{N}_0^N$ that specifies the integrator $z_{\theta,\lambda}(t) = \mathbb{E}(F(X_{\theta,\lambda}(t)))$ which satisfies (2.14). Due to Dynkin's Theorem [2] we can express the time-derivative of the integrator as

$$\dot{z}_{\theta,\lambda}(t) = \mathbb{E}(\mathbb{A}_{\theta,\lambda} F_{\theta,\lambda}(X_{\theta,\lambda}(t))).$$

Substituting this in (2.14) and observing that the resulting relation needs to hold for any initial state $X_{\theta,\lambda}(0) \in \mathcal{E}$ we can conclude that $F_{\theta,\lambda}$ must satisfy the *Poisson* equation (see Chapter 8 in [7])

$$\mathbb{A} F_{\theta,\lambda}(x) = \phi_{\text{out}}\left(\frac{\theta_1}{\theta_2}\right) - x_1 \qquad \text{for any} \qquad x \in \mathcal{E}. \tag{S26}$$

Such a function exists uniquely under our ergodicity assumption [8] and corresponding to this function $F_{\theta,\lambda}$ we define a bounded function $G_{\theta,\lambda}(x) = (G_{\theta,\lambda,1}(x), \dots, G_{\theta,\lambda,K-2}(x))$ by

$$G_{\theta,\lambda,k-2}(x) = \mathrm{Sign}\left(\Delta_k F_{\theta,\lambda}(x)\right) \quad \text{for} \quad k = 3, \dots, K, \tag{S27}$$

where the sign function is given by

$$\mathrm{Sign}(x) = \begin{cases} -1 & \text{if } x < 0 \\ 1 & \text{if } x \geq 0 \end{cases}$$

In other words, $G_{\theta,\lambda,k-2}(x)$ captures the signs of the actions of the difference operator $\Delta_k$ on the function $F_{\theta,\lambda}$ for the disturbance inducing reactions $k = 3, \dots, K$. We now restate Assumption 3.4 in the main text in a more formal way.

**Assumption S3.3** *For any $(\theta, \lambda) \in \Theta \times \Lambda_u$ we have the following:*

*(A)* **Stability:** *The state-space $\mathcal{E} = \mathbb{N}_0^N$ is irreducible for the CTMC dynamics $(X_{\theta,\lambda}(t))_{t\geq 0}$ and there exists a Foster-Lyapunov function $V_{\theta,\lambda}$ satisfying* (S5) *with $\mathbb{A} = \mathbb{A}_{\theta,\lambda}$. Hence this CTMC is exponentially ergodic with a unique stationary distribution $\pi_{\theta,\lambda}$. The projection map $f$ (see* (2.9)*) satisfies $\|f\|_{V_{\theta,\lambda}} < \infty$ where the norm $\|\cdot\|_{V_{\theta,\lambda}}$ is defined in* (S7)*.*

*(B)* **Local richness of the uncertain set $\Lambda_u$:** *Let $F_{\theta,\lambda}$ be the solution of the Poisson equation* (S26) *with $\mathbb{A} = \mathbb{A}_{\theta,\lambda}$ and let $G_{\theta,\lambda}(x)$ be the function given by* (S31)*. Then for any $\epsilon > 0$ there exist $R$ functions $\phi_1, \dots, \phi_R : \mathcal{E} \to \mathbb{R}_+$ such that*

$$\lambda_\gamma := \lambda + \sum_{i=1}^{R} \gamma_i \phi_i \in \Lambda_u \quad \text{for all} \quad \gamma = (\gamma_1, \dots, \gamma_R) \in (-\epsilon, \epsilon)^R \tag{S28}$$

*and*

$$\sum_{x \in \mathcal{E}} \left\| G_{\theta,\lambda}(x) - \sum_{i=1}^{R} c_i \phi_i(x) \right\|_{\ell_1} V_{\theta,\lambda}(x) \pi_{\theta,\lambda}(x) < \epsilon \tag{S29}$$

*for some constants $c_1, \dots, c_R$.*

The exponential ergodicity assumption is quite restrictive but it has been found that many biological networks readily satisfy this assumption [3] and this can be shown by constructing a linear Foster-Lyapunov function $V_{\theta,\lambda}$ (S6). Note that in the case of a linear $V_{\theta,\lambda}$, the condition $\|f\|_{V_{\theta,\lambda}} < \infty$ is trivially satisfied. From Theorem 4.5 in [4], we can see that if we define a measure $\mu$ over $\mathcal{E}$ as

$$\mu(x) = V_{\theta,\lambda}(x) \pi_{\theta,\lambda}(x)$$

then this is a finite measure i.e.

$$\sum_{x \in \mathcal{E}} \mu(x) < \infty.$$

Part (B) of Assumption 3.4 essentially says that the set $\Lambda_u$ of uncertain propensity maps is locally rich enough to allow perturbations by functions whose linear combination can closely approximate the vector-valued function $G_{\theta,\lambda}(x)$ in the sense of $\mathcal{L}_1$ norm over the measurable space $(\mathcal{E}, \mu)$. Before we prove our main result (Theorem 3.5) we need the following proposition.

**Proposition S3.4** *Suppose the conditions of Theorem 3.5 hold, and let $F_{\theta,\lambda}$ be the solution of the Poisson equation* (S26) *with $\mathbb{A} = \mathbb{A}_{\theta,\lambda}$ for some $(\theta, \lambda) \in \Theta \times \Lambda_u$. Then for the maxRPA property to hold, function $F_{\theta,\lambda}$ must be invariant under translations by the stoichiometric vectors $\zeta_k$ corresponding to disturbance inducing reactions i.e. for each $k = 3, \dots, K$ and each state $x \in \mathcal{E}$ such that $x \geq \nu_k$*

$$F_{\theta,\lambda}(x + \zeta_k) = F_{\theta,\lambda}(x). \tag{S30}$$

**Proof.** Fix a $(\theta, \lambda) \in \Theta \times \Lambda_u$ and an $\epsilon > 0$. Due to Assumption 3.4(B) there exist $R$ functions $\phi_1, \dots, \phi_R$ and constants $c_1, \dots, c_R$ such that (S29) holds. For any $\gamma = (\gamma_1, \dots, \gamma_R) \in (-\epsilon, \epsilon)^R$ let

$$\lambda_\gamma := \lambda + \sum_{i=1}^{R} \gamma_i \phi_i \in \Lambda_u.$$

Define a function $H_{\theta,\lambda}(x) = (H_{\theta,\lambda,1}(x), \dots, H_{\theta,\lambda,K-2}(x))$ by

$$H_{\theta,\lambda,k-2}(x) = \Delta_k F_{\theta,\lambda}(x) \quad \text{for} \quad k = 3, \dots, K, \tag{S31}$$

which implies that $G_{\theta,\lambda}(x) = \mathrm{Sign}(H_{\theta,\lambda}(x))$ where the sign function is being applied coordinate-wise. As $\|f\|_{V_{\theta,\lambda}} < \infty$ (see Assumption 3.4(A)), Theorem 2.3 in [8] implies that the same holds for the solution of the Poisson equation (S26) $F_{\theta,\lambda}(x)$. Hence there exists a constant $C_H > 0$ such that

$$\sup_{x \in \mathcal{E}} \max_{k=1,\dots,K-2} \frac{|H_{\theta,\lambda,k}(x)|}{V(x)} \leq C_H. \tag{S32}$$

From Theorem 3.3 in [9] it follows that for any $\gamma_i$ we have

$$\begin{aligned} \partial_{\gamma_i} \mathbb{E}_{\pi_{\theta,\lambda_\gamma}}(f)\Big|_{\gamma=\mathbf{0}} &= \sum_{k=3}^{K} \sum_{x \in \mathbb{N}_0^N} \partial_{\gamma_i} \lambda_k(x) \Delta_k F_{\theta,\lambda}(x) \pi_{\theta,\lambda}(x) \\ &= \sum_{k=1}^{K-2} \sum_{x \in \mathbb{N}_0^N} \phi_{ik}(x) H_{\theta,\lambda,k}(x) \pi_{\theta,\lambda}(x) \\ &= \sum_{x \in \mathbb{N}_0^N} \langle H_{\theta,\lambda}(x), \phi_i(x) \rangle \pi_{\theta,\lambda}(x), \end{aligned} \tag{S33}$$

where $\langle \cdot, \cdot \rangle$ denotes the standard inner product in $\mathbb{R}^{K-2}$, $f$ is the projection map (2.9) and $\phi_{ik}(x)$ is the $k$-th component of $\phi_i(x)$. For the maxRPA property to hold, the parametric sensitivity defined by the l.h.s. of (S33) must be 0 for each $\gamma_i$. Multiplying it by $c_i$ and summing over $i$ we get

$$0 = \sum_{x\in\mathbb{N}_0^N} \left\langle H_{\theta,\lambda}(x), \sum_{i=1}^{R} c_i\phi_i(x) \right\rangle \pi_{\theta,\lambda}(x)$$

$$= \sum_{x\in\mathbb{N}_0^N} \langle H_{\theta,\lambda}(x), G_{\theta,\lambda}(x)\rangle \pi_{\theta,\lambda}(x) - \sum_{x\in\mathbb{N}_0^N} \left\langle H_{\theta,\lambda}(x), G_{\theta,\lambda}(x) - \sum_{i=1}^{R} c_i\phi_i(x) \right\rangle \pi_{\theta,\lambda}(x).$$

Since

$$\langle H_{\theta,\lambda}(x), G_{\theta,\lambda}(x)\rangle = \sum_{k=1}^{K-1} |H_{\theta,\lambda,k-2}(x)| := \|H_{\theta,\lambda}(x)\|_{\ell_1}$$

we obtain

$$\sum_{x\in\mathbb{N}_0^N} \|H_{\theta,\lambda}(x)\|_{\ell_1}\pi_{\theta,\lambda}(x) = \sum_{x\in\mathbb{N}_0^N} \left\langle H_{\theta,\lambda}(x), G_{\theta,\lambda}(x) - \sum_{i=1}^{R} c_i\phi_i(x) \right\rangle \pi_{\theta,\lambda}(x)$$

$$\leq C_H \sum_{x\in\mathcal{E}} \left\| G_{\theta,\lambda}(x) - \sum_{i=1}^{R} c_i\phi_i(x) \right\|_{\ell_1} V_{\theta,\lambda}(x)\pi_{\theta,\lambda}(x)$$

$$\leq C_H\epsilon,$$

where $C_H$ is the constant in (S32) and the last inequality follows from (S29). As $\epsilon > 0$ is arbitrary, letting $\epsilon \to 0$ shows that

$$\sum_{x\in\mathbb{N}_0^N} \|H_{\theta,\lambda}(x)\|_{\ell_1}\pi_{\theta,\lambda}(x) = 0.$$

Due to irreducibility of the state-space $\mathcal{E} = \mathbb{N}_0^N$ and ergodicity of the CTMC (see Assumption 3.4(A)), we know that $\pi_{\theta,\lambda}(x) > 0$ for each $x \in \mathcal{E}$. This implies that $H_{\theta,\lambda}(x) = \mathbf{0}$ for each $x \in \mathbb{N}_0^N$ and completes the proof of this proposition. □

**Proof.**[Proof of Theorem 3.5] Applying the generator $\mathbb{A}_{\theta,\lambda}$ to the function $F_{\theta,\lambda}$ and using the translation invariance shown by Proposition S3.4, we obtain

$$\mathbb{A}_{\theta,\lambda}F_{\theta,\lambda}(x) = \theta_1 m_1(x)(F_{\theta,\lambda}(x+\zeta_1) - F_{\theta,\lambda}(x)) + \theta_2 m_2(x)(F_{\theta,\lambda}(x+\zeta_2) - F_{\theta,\lambda}(x)).$$

Since $F_{\theta,\lambda}$ solves the Poisson equation (S26), the r.h.s. of this equation must equate to the r.h.s. of (S26) for all $x \in \mathbb{N}_0^N$. This can only happen if for some distinct $j_1, j_2 \in \{1,2\}$

$$m_{j_1}(x) \equiv 1 \quad \text{and} \quad F_{\theta,\lambda}(x+\zeta_{j_1}) - F_{\theta,\lambda}(x) = \frac{1}{\theta_{j_1}}\phi_{\text{out}}\left(\frac{\theta_1}{\theta_2}\right) \tag{S34}$$

and either

$$m_{j_2}(x) = x_1 \quad \text{and} \quad F_{\theta,\lambda}(x+\zeta_{j_2}) - F_{\theta,\lambda}(x) = -\frac{1}{\theta_{j_2}} \tag{S35}$$

or

$$m_{j_2}(x) = 1 \quad \text{and} \quad F_{\theta,\lambda}(x+\zeta_{j_2}) - F_{\theta,\lambda}(x) = -\frac{x_1}{\theta_{j_2}}. \tag{S36}$$

We shall later argue that condition (S36) cannot be satisfied at all $x \in \mathbb{N}_0^N$. Hence we proceed under the assumption that (S35) holds. Without loss of generality we can assume that $j_1 = 1$ and $j_2 = 2$ (the other case is symmetric) and $F_{\theta,\lambda}(\mathbf{0}) = 0$ where $\mathbf{0}$ is the $N$-dimensional vector of zeros. Observe that the form of the mass-action monomials $m_1(x)$ and $m_2(x)$ in (S34) and (S35) implies part (A).

Applying the parameter sensitivity formula given by Theorem 3.3 in [9] we can conclude that

$$\partial_{\theta_1}\mathbb{E}_{\pi_{\theta,\lambda}}(f) = \sum_{x\in\mathbb{N}_0^N} m_1(x)(F_{\theta,\lambda}(x+\zeta_1) - F_{\theta,\lambda}(x))\pi_{\theta,\lambda}(x) = \frac{1}{\theta_2}\phi'_{\text{out}}\left(\frac{\theta_1}{\theta_2}\right) \tag{S37}$$

$$\text{and} \quad \partial_{\theta_2}\mathbb{E}_{\pi_{\theta,\lambda}}(f) = \sum_{x\in\mathbb{N}_0^N} m_2(x)(F_{\theta,\lambda}(x+\zeta_2) - F_{\theta,\lambda}(x))\pi_{\theta,\lambda}(x) = -\frac{\theta_1}{\theta_2^2}\phi'_{\text{out}}\left(\frac{\theta_1}{\theta_2}\right). \tag{S38}$$

However (S34) and (S35) imply that

$$\sum_{x\in\mathbb{N}_0^N} m_1(x)(F_{\theta,\lambda}(x+\zeta_1) - F_{\theta,\lambda}(x))\pi_{\theta,\lambda}(x) = \frac{1}{\theta_1}\phi_{\text{out}}\left(\frac{\theta_1}{\theta_2}\right) \tag{S39}$$

$$\text{and} \quad \sum_{x\in\mathbb{N}_0^N} m_2(x)(F_{\theta,\lambda}(x+\zeta_2) - F_{\theta,\lambda}(x))\pi_{\theta,\lambda}(x) = -\frac{1}{\theta_2}\phi_{\text{out}}\left(\frac{\theta_1}{\theta_2}\right). \tag{S40}$$

Equating (S37) and (S39) we get

$$\phi_{\text{out}}\left(\frac{\theta_1}{\theta_2}\right) = \frac{\theta_1}{\theta_2}\phi'_{\text{out}}\left(\frac{\theta_1}{\theta_2}\right)$$

which also ensures that (S38) and (S40) are equal. This relation proves that

$$\phi_{\text{out}}(x) = \kappa x \tag{S41}$$

for some $\kappa > 0$. We shall soon see this this $\kappa$ is exactly as stated in part (B) of this theorem.

Let $e_m$ be the $m$-th standard basis vector in $\mathbb{R}^N$ (i.e. its $m$-th component is one and the rest are all zeros). As the state-space $\mathbb{N}_0^N$ is irreducible for the dynamics, for each species $\mathbf{X}_i$ there exists a sequence of reactions $k_1^{(i)}, \dots, k_n^{(i)}$ such that there is a positive probability of going from state $\mathbf{0}$ to state $e_i$ via these reactions. Due to (2.5) we must have

$$z_j := \sum_{l=1}^{j} \zeta_{k_l^{(i)}} \geq \nu_{k_{j+1}^{(i)}} \quad \text{for each} \quad j = 0, 1, \dots, n-1$$

and

$$z_n := \sum_{l=1}^{n} \zeta_{k_l^{(i)}} = e_i.$$

We can also observe that for any $z \in \mathbb{N}_0^N$ this same sequence of reactions will also allow the dynamics to go from state $z$ to state $(z+e_i)$ with a positive probability (see [5]). Setting $z_0 = \mathbf{0}$ we note that

$$\begin{aligned} F_{\theta,\lambda}(z+e_i) - F_{\theta,\lambda}(z) &= \sum_{j=1}^{n} \left( F_{\theta,\lambda}(z+z_j) - F_{\theta,\lambda}(z+z_{j-1}) \right) \\ &= \sum_{j=1}^{n} \left( F_{\theta,\lambda}(z+z_{j-1}+\zeta_{k_j^{(i)}}) - F_{\theta,\lambda}(z+z_{j-1}) \right). \end{aligned}$$

Observe that because of conditions (S30), (S34) and (S35) we obtain

$$\theta_2(F_{\theta,\lambda}(z+z_{j-1}+\zeta_{k_j^{(i)}}) - F_{\theta,\lambda}(z+z_{j-1})) = \begin{cases} 0 & \text{if } k_j^{(i)} \in \{3,\dots,K\} \\ \kappa & \text{if } k_j^{(i)} = 1 \\ -1 & \text{if } k_j^{(i)} = 2. \end{cases}$$

Therefore if $n_i^+$ and $n_i^-$ denote the number of instances of reactions 1 and 2 respectively. Then we have

$$\theta_2(F_{\theta,\lambda}(z+e_i) - F_{\theta,\lambda}(z)) = q_i := \kappa n_i^+ - n_i^- \tag{S42}$$

for any $z \in \mathbb{N}_0^N$. This allows us to identify an integer $q_i$ with each species $\mathbf{X}_i$ and it shows that $F_{\theta,\lambda}$ must be of the form (3.29). Proposition S3.4 implies that $q$ must be orthogonal to the stoichiometric vectors of the disturbance inducing reactions, i.e.

$$\langle q, \zeta_k \rangle = 0 \quad \text{for} \quad k = 3,\dots,K. \tag{S43}$$

Moreover since (S41) holds, (S34) and (S35) imply that

$$\langle q, \zeta_1 \rangle = \kappa \qquad \text{and} \qquad \langle q, \zeta_2 \rangle = -1. \tag{S44}$$

Taken together, (S3.2) and (S44), prove that the vector $q = (q_1,\dots,q_N)$ must satisfy the linear system (3.17).

We now argue that under our assumptions, (S36) cannot hold at each $x \in \mathbb{N}_0^N$. Suppose that this condition indeed holds. Then what changes in the preceding analysis is that now

$$\theta_2(F_{\theta,\lambda}(z+z_{j-1}+\zeta_{k_j^{(i)}}) - F_{\theta,\lambda}(z+z_{j-1})) = -f(z+z_{j-1}) \quad \text{if} \quad k_j^{(i)} = 2,$$

where $f$ is the projection map (2.9) that maps the state vector to the state of the output species. Note that in this case the l.h.s. of (S42) will depend on the choice of the reaction path, and the intermediate states encountered, in going from state $z$ to state $(z+e_i)$. This violates the fact that the solution $F_{\theta,\lambda}$ of the Poisson equation (S26) is unique up to addition by a constant.

So far we had assumed that $j_1 = 1$ and $j_2 = 2$ in (S34) and (S35). The treatment of the other symmetric case ($j_1 = 2$ and $j_2 = 1$) is similar to above. In this case $\phi_{\text{out}}(x) = (\kappa x)^{-1}$. This proves the "only if" part of the result.

The "if" part of the result follows from ergodicity of the stochastic dynamics along with the presence of an integrator (3.29) that satisfies (2.14). This completes the proof of this theorem. □

#### S3.3 Generic decomposition of stochastic maxRPA networks

In this section we study the structure of a maxRPA network in the stochastic setting. We restrict ourselves with *bimolecular* reactions in which all reactions can have at most two reactants i.e. $\sum_i \nu_{ik} \leq 2$ for each reaction $k$ of the form (2.1). We shall show that in the stochastic setting, a maxRPA network cannot be homothetic but *it needs* to be antithetic. A key property of such antithetic networks is the existence of a generalised *annihilation* or *sequestration* reaction between the internal model (IM) (comprising species in $\mathcal{C}_\pm$) and the *rest of the network* (RON) (comprising species in $\mathcal{C}_0$). This reaction is of the form

$$\mathbf{X}_p + \mathbf{X}_\ell \to \mathcal{C}_0^*, \tag{S45}$$

where $\mathbf{X}_p \in \mathcal{C}_+$, $\mathbf{X}_\ell \in \mathcal{C}_-$ and $\mathcal{C}_0^*$ denotes any combination of species that belong to the set $\mathcal{C}_0$. Note that such a reaction cannot arise in a homothetic network because either $\mathcal{C}_+$ or $\mathcal{C}_-$ is empty.

**Proof.**[Proof of Theorem 3.8] We first prove by contradiction that no maxRPA network in the stochastic setting can be homothetic (recall Definition 3.7). Consider a maxRPA network in the stochastic setting characterised by a pair $(q, \kappa)$ that satisfies the linear-algebraic system (3.17) (see Theorem 3.5(B)). Suppose that $\mathcal{C}_- = \emptyset$ and hence all nonzero components are $q$ are positive. From Theorem 3.5(A) we know that the second reaction must be of the form $\mathbf{X}_1 \to *$ and its stoichiometry vector $\zeta_2$ satisfies

$$q^T \zeta_2 = -1. \tag{S46}$$

Note that except for the first coordinate (corresponding to the output species $\mathbf{X}_1$) all other components of $\zeta_2$ must be nonnegative. Since $q_1 = 0$ and all other components of $q$ are nonnegative, we see that (S46) cannot hold, resulting in a contradiction. Now we consider the other case that $\mathcal{C}_+ = \emptyset$ and so all nonzero components are $q$ are negative. Again from Theorem 3.5(A) we know that the first reaction must be of the form $\emptyset \to *$ and its stoichiometry vector $\zeta_1$ satisfies

$$q^T \zeta_1 = \kappa > 0. \tag{S47}$$

As all components of $\zeta_1$ need to be nonnegative and all components of $q$ are nonpositive, (S47) yields a contradiction. This shows that the maxRPA network cannot be homothetic and proves part (A) of this theorem.

We now prove part (B) with the assumption that the maxRPA network is antithetic and hence $q$ has both positive and negative components. As all components of $\zeta_1$ are nonnegative, condition (S47) implies that there exists some species, let us call it $\mathbf{X}_2$, with $q_2 > 0$ and $\zeta_{21} > 0$[1]. Hence $\mathbf{X}_2 \in \mathcal{C}_+$ and the first reaction is of the form (3.32). This completes the proof of part (B). The proof of part (C) is similar. Lemma S3.2 shows that $q_1 = 0$, and so from (S46) we can conclude that there exists some species, let us call it $\mathbf{X}_3$, with $q_3 < 0$ and $\zeta_{32} > 0$. Hence $\mathbf{X}_3 \in \mathcal{C}_-$ and the second reaction is of the form (3.33). This completes the proof of part (C).

We now prove part (D). Let $I_0$ be the set of addresses of species in $\mathcal{C}_0$ respectively, i.e.

$$I_0 = \{i = 1, \dots, N : \mathbf{X}_i \in \mathcal{C}_0\}.$$

[1] Here $\zeta_{21}$ denotes the second component of the first stoichiometric vector $\zeta_1$

Define $\mathcal{S}_0 \subset \mathbb{N}_0^N$ to be the set given by

$$\mathcal{S}_0 = \{x = (x_1, \dots, x_N) \in \mathbb{N}_0^N : x_i = 0 \quad \text{for each} \quad i \notin I_0\}$$

and so if the state is in $\mathcal{S}_0$ then all the species in the internal model (i.e. in the set $\mathcal{C}_\pm$) have zero copy-numbers. Pick any species $\mathbf{X}_m \in \mathcal{C}_\pm$. Since the state-space $\mathcal{E} = \mathbb{N}_0^N$ is irreducible, there exists a sequence of reactions $k_1, \dots, k_n$ such that there is a positive probability of going from state $e_m$ to state $\mathbf{0}$ via these reactions, thereby eliminating exactly one molecule of species $\mathbf{X}_m$. For this to hold we must have

$$z_j := e_m + \sum_{l=1}^{j} \zeta_{k_l} \geq \nu_{k_{j+1}} \quad \text{for each} \quad j = 0, 1, \dots, n-1$$

and

$$z_n := e_m + \sum_{l=1}^{n} \zeta_{k_l} = \mathbf{0}.$$

Note that $z_n \in \mathcal{S}_0$ and $z_0 = e_m \notin \mathcal{S}_0$. Let $k_*$ be the first reaction in this sequence which originates from a state outside $\mathcal{S}_0$ and leads to a state within $\mathcal{S}_0$, i.e. $k_* = k_l$ where

$$l = \min\{j = 1, \dots, n : z_{j-1} \notin \mathcal{S}_0 \quad \text{and} \quad z_j \in \mathcal{S}_0\}.$$

Observe that $k_*$ needs to eliminate molecules of some species $\mathbf{X}_p$ in $\mathcal{C}_\pm$ and all its products must be species in $\mathcal{C}_0$. Hence this reaction $k_*$ cannot be the first reaction (3.32) or the second reaction (3.33). Since all reactions are bimolecular this reaction $k_*$ must necessarily be of the form

$$\mathbf{X}_p \longrightarrow \mathcal{C}_0^* \tag{S48}$$

or of the form

$$\mathbf{X}_p + \mathbf{X}_\ell \longrightarrow \mathcal{C}_0^*. \tag{S49}$$

As $q$ satisfies (3.17), reaction $k_*$ being of the form (S48) would imply that $q_p = 0$ which is a contradiction since $\mathbf{X}_p$ is in $\mathcal{C}_\pm$. Therefore reaction $k_*$ must be of the form (S49) and in this case condition (3.17) implies that $q_p + q_\ell = 0$ or

$$q_p = -q_\ell.$$

Clearly for this to happen one of the species $\mathbf{X}_p$ or $\mathbf{X}_\ell$ belongs to $\mathcal{C}_+$ and the other belongs to $\mathcal{C}_-$, thereby proving the existence of a generalised antithetic reaction (3.30). This analysis also shows that all reactions where the products are species in $\mathcal{C}_0$ and one of the reactants is a species in $\mathcal{C}_\pm$, must necessarily be of the form (3.30). This completes the proof of this theorem. □

**Remark S3.5** *Notice that the proof of part (D) of Theorem 3.8 only hinges on the assumption that the state-space $\mathbb{N}_0^N$ is irreducible for the reaction network (see Assumption*

*3.4(A)). As we mention in Section 3.2 of the main text, this irreducibility assumption typically holds when the stoichiometry matrix satisfies the full-rank condition* (3.16) *(see [5] for more details). Hence for most bimolecular deterministic antithetic maxRPA networks satisfying* (3.31)*, the assertion of part (D) of this theorem continues to hold, and in particular such networks would have a generalised sequestration reaction of the form* (3.30)*. If the deterministic maxRPA network is homothetic, this argument cannot be applied as for these networks the state-space* $\mathbb{N}_0^N$ *will not be irreducible for the corresponding stochastic network.*
